# Gut bacteria regulate the pathogenesis of Huntington’s disease in *Drosophila* model

**DOI:** 10.1101/2021.08.12.456124

**Authors:** Anjalika Chongtham, Jung Hyun Yoo, Theodore M. Chin, Ngozi D. Akingbesote, Ainul Huda, Ali Khoshnan

## Abstract

Changes in the composition of gut microbiota are implicated in the pathogenesis of several neurodegenerative disorders. Here, we investigated whether gut bacteria affect the progression of Huntington’s disease (HD) in transgenic *Drosophila melanogaster* (fruit fly) models expressing human full-length or N-terminal fragments of mutant huntingtin (HTT) protein, here referred to as HD flies. We find that elimination of commensal gut bacteria by antibiotics reduces the aggregation of amyloidogenic N-terminal fragments of HTT and delays the development of motor defects. Conversely, colonization of HD flies with *Escherichia coli* (*E. coli*), a known pathobiont of human gut with links to neurodegeneration, accelerates HTT aggregation, aggravates immobility and shortens lifespan. Similar to antibiotics, treatment of HD flies with small compounds such as luteolin, a flavone, or crocin a beta-carotenoid, ameliorates disease phenotypes and promotes survival. Crocin prevents colonization of *E. coli* in the gut and alters the abundance of commensal bacteria, which may be linked to its protective effects. The opposing effects of *E. coli* and crocin on HTT aggregation, motor defects and survival in transgenic *Drosophila* models support the involvement of gut-brain networks in the pathogenesis of HD.

## 1. Introduction

Huntington’s disease (HD) is a progressive genetically inherited neurodegenerative disorder characterized by debilitating motor, psychiatric, and cognitive symptoms (Bates et al., 2015, Ghosh and Tabrizi, 2018). Expansion of a CAG repeat (>35) in the exon 1 of huntingtin (HTT) gene, which translates into an abnormal polyglutamine (polyQ) tract, is the underlying cause of HD (Huntington’s Disease Collaborative Research Group, 1993). The expanded polyQ enhances the amyloidogenic properties of HTT exon1 (HTTex1) peptide, which misfolds and forms insoluble protein assemblies in neurons (Difiglia et al., 1997). Expansion of polyQ repeat is a major determinant of disease onset in HD, however, multiple genetic and environmental factors may affect the development and progression of symptoms. One potential modifier is neuroinflammation exemplified by elevated levels of inflammatory microglia and TH17.1 cells in the brains of pre-manifest HD patients. Increase in activated immune cells coincides with elevated production of inflammatory cytokines, which persists during the symptomatic stages of HD (Björkqvist et al., 2008, Politis et al., 2015, Von Essen et al., 2020). Although mutant HTT has directly been implicated in the activation of inflammatory pathways, the environmental inducers of inflammation in HD remain unknown (Khoshnan et al., 2004, Trager et al., 2014, Khoshnan et al., 2017).

Commensal and acquired gut microorganisms are prominent sources of inflammogens implicated in neurological disorders. Within the last decade, extensive studies demonstrate that the gastrointestinal (GI) tract and its resident microbes (collectively known as microbiota, and their genomes as microbiome) regulate the nervous system physiology. For example, gut microbiota influences neurodevelopment, neurodegeneration, neurotrophin and neurotransmitter production, neuropsychiatric and motor behaviors, and neuroinflammation (Sharon and Mazmanian, 2016, Sampson et al., 2016, Morais et al., 2021). Changes in the homeostasis of gut microbiota (dysbiosis) have been linked to the pathogenesis of several neuropsychiatric and neurodegenerative disorders, including, autism spectrum disorder (ASD), multiple sclerosis (MS), Alzheimer’s disease (AD), Parkinson’s disease (PD) and amyotrophic lateral sclerosis (ALS) (Hsiao et al., 2013, Sampson et al., 2016, Marizzoni et al., 2020, Takewaki et al., 2020, Zeng et al., 2020). Neuroinflammation is a prominent feature of gut dysbiosis in neurodegenerative disorders, which includes the activation of microglia and subsequent production of inflammatory cytokines (Sampson et al., 2016, Abdel-Haq et al., 2018). Notably, inflammatory bacteria such as *Enterobacteriaceae* are elevated in the gut of PD patients and their abundance correlates with worsening of the symptoms and pathology (Keshavarzian et al., 2015, Scheperjans et al., 2015, Li et al., 2017). Moreover, lipopolysaccharides (LPS), major components of gram negative bacteria, which include *Enterobacteriaceae,* have been implicated in the pathogenesis of PD (Perez-Pardo et al., 2019). Gut dysbiosis manifested by reduced alpha diversity in the bacterial communities were recently reported in a cohort of HD patients. Notably, in these studies changes in the abundance of *Eubacterium halii* and potentially other candidates coincide with altered cognition (Wasser et al., 2020). Another gut microbiome analysis of a group of HDs patient’s links blooming of the gram-negative bacteria *Bilophila* species to elevated levels of inflammatory cytokines (Du et al., 2021). In transgenic mouse models of HD expressing the neurotoxic mutant HTTex1, gut dysbiosis has been linked to weight loss, motor deficit, metabolic changes and disruption in the intestinal epithelium (Kong et al., 2020, Stan et al., 2020, Kong et al., 2021). These encouraging studies highlight a potential role of gut microbiota in the pathogenesis of HD. However, the mechanisms of how a specific bacterium may affect disease manifestation remain to be investigated.

*Drosophila melanogaster* (fruit fly) is emerging as a useful model to study the impact of gut-brain interactions in the development and progression of neurological disorders (Wu et al., 2017, Douglas AE, 2018). Similar to mammals, gut homeostasis in *Drosophila* is regulated by the interaction of bacteria with the enteric neurons and intestinal epithelium including the enteroendocrine cells, which produce antimicrobial peptides (AMPs) involved in immunity against invasive pathogens and maintaining optimal abundance of commensal bacteria (Hanson and Lemaitre, 2020). *Drosophila* has a handful of commensal gut bacteria, which can easily be manipulated for studies on gut-brain communications (Douglas AE, 2018, Ankrah et al., 2021). *Lactobacillus* and *Acetobacter* species are the two most abundant bacterial genera, which regulate nutrients acquisition, growth, metabolism, immune development and locomotion (Schretter et al., 2018, Storelli et al., 2018, Henriques et al., 2020, Yamauchi et al., 2020, Ankrah et al., 20201). The *Drosophila* models of HD display several hallmarks of disease progression including protein aggregation, motor defects, and aberrant expression of genes including those implicated in systemic inflammation (Barbaro et al., 2015, Al-Ramahi et al., 2018). The simplicity of *Drosophila* gut microbiota offers a useful platform to rapidly examine the influence of endogenous and single exogenous bacterial species on HD pathogenesis and to investigate the role of HTT in gut–brain pathways. As a first step, we explored whether manipulation of gut bacteria in HD *Drosophila* models or colonization with human pathobiont *E. coli* influences disease development. Here, we report that gut bacteria promote the aggregation of amyloidogenic N-terminal fragments of HTT, contribute to development of aberrant motor behavior and reduce the life span of HD flies. We further provide evidence that editing the gut environment of HD flies is a potential therapeutic target.

## 2. Materials and methods

### 2.1. Fly stocks

The huntingtin expressing transgenic *Drosophila* lines used in this study were M{UAS-HTT.ex1.Q25}, M{UAS-HTT.ex1.Q120}, M{UAS-HTT.FL.Q25}, M{UAS-hHTT.fl.Q120}and M{UAS-HTT.586.Q120} (Barbaro et al., 2015, Chongtham et al., 2020). The GAL4 drivers used in the experiments were the pan-neuronal *elav*-Gal4 C155 driver (Bloomington Stock Number B#458) and ubiquitous *da*-Gal4 (B#8641). Fly cultures were maintained on standard cornmeal/sugar/agar media on a 12:12 hour light:dark cycle. Appropriate crosses were carried out to obtain desired progenies. Briefly, standard food vials with 10 males and 10 females of appropriate genotype were allowed to mate for 2 days and then passed into new vials at 20°C. All assays used female progeny, which were collected after the eclosion and maintained at 20°C for 2-3 days until enough flies had been collected.

### 2.2. Treatment of flies with antibiotics or small molecules

For drug treatment, groups of 15-20 female flies were placed in vials containing standard cornmeal/sugar/agar food alone for non-treatment control experiments or standard food mixed with 1mg/ml of water-soluble test compounds crocin (cat# 17304 Sigma), rifaximin (cat# Y0001074, Sigma), luteolin (# L9283 Sigma) or 1% penicillin-streptomycin (cat#15140-122; Gibco). The flies were transferred to 25°C and passaged to fresh vials every second or third day. For treatment with live bacteria, curli producing (MC4100) and curli deficient (Δcsg) *E. coli* strains from −80°C frozen stocks were grown on YESCA (1% Casamino Acids, 0.12% yeast extract, 2% Bacto agar) agar plates at 25°C for 48 hr. *Lactobacillus rhamnosus* strain (JB-1, ATCC) from frozen stock was streaked on MRS (ingredient) agar plates and grown at 37°C overnight. *Acetobacter* stock was prepared by grinding adult flies in PBS followed by culturing of isolated bacteria in mannitol agar plate. Isolated colonies were grown in liquid manitol broth at 37°C in bacterial shaker. The bacterial cultures were harvested by centrifugation (3000xg, 5min), washed and resuspended in PBS. The bacterial suspension (0.5 OD or 5×10^7^ cells) was then mixed with standard food and groups of 15-20 female progeny which had been first treated with 1% penicillin-streptomycin-supplemented food for 3 to 4 days, were transferred to the bacteria supplemented food. The flies were transferred to new food with the live bacteria every second or third day at 25°C.

To examine the effects of crocin on *E. coli* treated flies, crocin (1mg/mL) and *E.coli* (0.5OD or 5×10^7^ cells) were both mixed with standard food and flies, which had been treated with 1% penicillin-streptomycin for 3-4 days, were transferred to the crocin and *E. coli* mixed food. Flies were transferred to fresh food every second or third day and their bacterial load and motor behavior was monitored.

### 2.3. Western blotting

At least 10 wandering third instar larvae or adult flies were lysed and homogenized in RIPA buffer (25mM Tris-HCl pH7.6, 150mM NaCl, 1% NP-40, 1mM EDTA) with protease inhibitors (Complete, Mini Protease Inhibitor Cocktail, Roche Applied Science). The lysates were then boiled at 95°C for 5 min, and equal amount of proteins were separated by SDS/PAGE on pre-cast 4-20% polyacrylamide gradient gels (Cat# 5671094, Biorad) and transferred to immune-blot PVDF membrane (Merck cat# IPVH00010). Membranes were blocked with blocking solution (5% non-fat milk in 0.05% Tween in PBS) and incubated with primary anti-HTT antibody PHP1 (1:1000 in blocking solution) overnight at 4°C. The blots were then treated with HRP-conjugated goat anti-moue secondary antibody (1:10000) diluted in blocking solution for 1 h and developed with enhanced chemiluminescent (ECL) substrate (Cat#1705060, Biorad).

### 2.4. SDD-AGE

To separate SDS-resistant amyloid assemblies, semi-denaturing detergent agarose gel electrophoresis (SDD-AGE) was performed (Halfmann and Lindquist, 2008) with some modifications. Fly lysates were prepared as described above and resolved by electrophoresis in SDD-AGE gels (1.5% agarose, 1X TAE, 0.1% SDS). Proteins were transferred to immune-blot PVDF membrane by overnight downward capillary action using 1X TBS. Membranes were then treated as western blots above.

### 2.5. Immunostaining of larval brains and guts of adult flies

Wandering third instar larvae were cut into anterior and posterior halves, and the anterior halves were turned inside out and placed in PBS on ice. These halves were then fixed by rocking for 30 min at RT with 4 % formaldehyde made in PBST (PBS + 0.2 % Triton X-100). After fixation, halves were washed three times with PBT, blocked with 5 % BSA in PBT for 1 hr at RT, probed overnight with primary antibody (s) at 4°C, washed, blocked again and incubated with secondary antibody (s) for 2 hr and washed again. Larval brains were then dissected out and mounted in Vectashield-DAPI medium. The primary antibodies were rat-Elav-7E8A10 anti-elav (used at 1:200 dilution in PBS; Developmental Studies Hybridoma Bank), PHP1 anti-HTT (used at 1:500 dilution in PBS; Ko et al., 2018). The secondary antibodies were Alexa Fluor 488 goat anti-mouse (green) and Alexa Fluor 568 goat anti-rat (red) (used at 1:250 dilution in blocking solution, Life Technologies).

Whole guts of adult flies were dissected out in PBS, fixed in 4% formaldehyde in PBT for 1 h, washed with PBT and incubated with blocking buffer for 1h at RT. The guts were then probed with anti-*E. coli* monoclonal antibody (produced in house) overnight at 4°C. After washing with PBST, guts were incubated with Alexa Fluor 488 goat anti-mouse antibody diluted in blocking buffer for 2 hr at RT and washed. The guts were mounted in Vectashield-DAPI medium. Images of mounted tissues were captured using a Leica Sp8 laser scanning microscope and analyzed using Leica Application Suite X (LAS X) software.

### 2.6. Climbing Assay

For monitoring the locomotor ability, 10 flies were gently tapped to the bottom of a vertical glass vial (diameter, 2.2 cm)., as adapted from Liu et al., 2008. The number of flies that climbed a height of 5 cm within 10 s was recorded. The test was repeated three times each for a group of 10 flies at 1 min intervals.

### 2.7. Longevity assay

A longevity assay was performed as described previously with slight modifications (Barbaro et al., 2015). Briefly, eighty eclosed female flies (1-3 days old) were divided into groups 10 with the indicated treatment in the figure legends and were maintained at 20℃ with a 12:12 hr light:dark cycle. Longevity assays in the presence of *E.coli* were performed at 25℃ to accommodate bacterial growth. Progenies were and transferred to fresh vials with standard food +/- treatment. Cultures were monitored every other day and the number of dead flies counted.

### 2.8. Congo Red staining of bacterial colonies and CFU counting

To determine the *E. coli* (Curli-producing) load in the gut, flies were surface sterilized with 70% ethanol for 1 min and washed three times with sterile 1X PBS. Flies were then homogenized in groups of 5 in 500 μL of sterile 1X PBS using a motorized pestle. Microbial counts were determined by serial dilution plating of the homogenates on YESCA agar plates supplemented with 50 µg/mL of Congo Red (CR) (Sigma). The YESCA CR plates were incubated at 25°C for 2 days to induce curli production and the number of red colonies that showed curli expression were counted.

### 2.9. Quantification of dead pupae

Five groups of ten UAS-HTTex1-120Q males and elav-Gal4 female virgins were allowed to mate for 2 days at 20℃ and transferred into vials with standard food supplemented with crocin (1mg/mL) or 1% penicillin-streptomycin for laying eggs. Twenty days after crosses were made, all flies were emptied from the vials, and empty pupal cases and cases containing dead flies were counted to calculate the percentage of pupal survival (Hatfield et al., 2015).

### 2.10. 16S sequencing of gut bacteria

Flies were harvested on the indicated days post-eclosion, gently disinfected in 70% ethanol and 3 subsequent rinse in PBS to remove any external bacteria and stored immediately at −80°C. Similar samples for different batches of flies were mixed and shipped to sequencing facility at Zymo Research Irvine, CA. Briefly, bacterial DNA was extracted using ZymoBIOMICS-96 MagBead DNA kit (Zymo Research Irvine, CA). The DNA samples were prepared for targeted sequencing with the Quick-16S NGS library Prep kit and 16S primer set V3-V4 (Zymo Research Irvine, CA). The final library was sequenced on Ilumina MiSeq with a V3 reagent Kit (600 cycles). The sequencing was performed with 10% PhiX Spike-in.

### 2.11. Bioinformatics Analysis of bacteria

Unique amplicon sequences were inferred from raw reads using the Dada2 pipeline (Callahan et al., 2016). Chimeric sequences were also removed with the Dada2 pipeline. Taxonomy assignment was performed using Uclust from Qiime v.1.9.1. Taxonomy was assigned with the Zymo Research Database, a 16S database that is internally designed and curated, as reference. Composition visualization, alpha-diversity, and beta-diversity analyses were performed with Qiime v.1.9.1 (Caporaso et al., 2010). If applicable, taxonomy that have significant abundance among different groups were identified by LEfSe (Segata et al., 2011) using default settings. Other analyses such as heatmaps, Taxa2SV_deomposer, and PCoA plots were performed with internal scripts.

### 2.12. Absolute Abundance Quantification

A quantitative real-time PCR was set up with a standard curve. The standard curve was made with plasmid DNA containing one copy of the 16S gene and one copy of the fungal ITS2 region prepared in 10-fold serial dilutions. The primers used were the same as those used in Targeted Library Preparation. The equation generated by the plasmid DNA standard curve was used to calculate the number of gene copies in the reaction for each sample. The PCR input volume (2 μl) was used to calculate the number of gene copies per microliter in each DNA sample. The number of genome copies per microliter DNA sample was calculated by dividing the gene copy number by an assumed number of gene copies per genome. The value used for 16S copies per genome is 4. The value used for ITS copies per genome is 200. The amount of DNA per microliter DNA sample was calculated using an assumed genome size of 4.64 x 106 bp, the genome size of *Escherichia coli,* for 16S samples, or an assumed genome size of 1.20 x 107 bp, the genome size of *Saccharomyces cerevisiae*, for ITS samples. This calculation is shown below: Calculated Total DNA = Calculated Total Genome Copies × Assumed Genome Size (4.64 × 106 bp) × Average Molecular Weight of a DNA bp (660 g/mole/bp) ÷ Avogadro’s Number (6.022 x 1023/mole).

### 2.13. Statistical analysis

Error bars show Standard Error of the Mean (SEM = standard deviation/square root of n). Statistical significance was established using Student’s t-test for pairwise and analysis of variance (ANOVA) with Tukey’s post hoc test for multiple comparisons on Prism software (GraphPad) (*=P<.05, **=P<.01, ***=P<.001).

## 3. Results

### 3.1. Gut Bacteria promote the aggregation of amyloidogenic fragments of HTT in Drosophila *models of HD*

Transgenic *Drosophila* (fruit fly) expressing HTTex1 (120Qs) under the control of pan-neuronal driver elav-Gal4 (Ex1-HD) displays gut dysbiosis exemplified by elevated levels of total bacteria when compared to a line expressing WT HTTex1 (25Qs) (Supplementary fig. 1A-C). Thus, a toxic function of neuronal mutant HTTex1 may be disruptions in gut homeostasis. To examine whether gut bacteria regulate the aggregation of HTTex1, we generated larvae of Ex1-HD line in the presence of a gut-specific antibiotic rifaximin to eliminate bacteria (Supplementary fig. 1D) and examined for the accumulation of HTTex1 assemblies by Western blots (WBs) and immunohistochemistry (IHC). Rifaximin treatment significantly reduces the levels of HTTex1 aggregates in the nervous system of Ex1-HD larvae (Fig. 1A and B). Given that HTTex1 is selectively expressed in neurons, these findings suggest that gut bacteria may alter neuronal physiology and pathways, which regulate the misfolding and aggregation of HTTex1. We then asked whether gut bacteria with links to neurodegeneration in humans may have a similar effect. *E. coli* a gut pathobiont, has been implicated in the pathogenesis of PD. In particular, colonization of transgenic mice expressing human α-synuclein or normal rats with an *E. coli* strain expressing the functional bacterial amyloids curli accelerates the aggregation of α-synuclein in the brain, induces neuroinflammation and worsens motor symptoms (Chen et al., 2016, Sampson et al., 2020). We used an HD fly line ubiquitously expressing the N-terminal 586 AA fragment of mutant HTT (120Qs) (N-586 HD) to explore the impact of *E. coli* on aggregation. Protein aggregates in this line accumulate much slower than in the Ex1-HD line thus, making it ideal for these assays (Barbaro et al., 2015). We generated the N-586 HD larvae on food containing an *E. coli* strain expressing curli or an isogenic mutant where the operon for curli has been deleted (Reichhardt et al., 2015). We find that colonization with either strain accelerates the aggregation of N-586 HTT fragment (Fig. 1C). While we do not observe any selective effects of curli on HTT proteostasis in the N-586 HD flies, recombinant curli promotes the aggregation of mutant HTTex1-EGFP when co-expressed in HEK-293 tissue culture cells (Supplementary fig. 2). A likely explanation is direct interaction of curli with mutant HTT may be essential to induce misfolding, whereas other dominant aggregation-promoting components of *E. coli* common to both strains may function by indirect pathways. Collectively, these findings identify gut bacteria as promoters of HTT aggregation in fruit fly models of HD. We also explored whether *E. coli* induces protein aggregation in transgenic flies ubiquitously expressing full-length mutant HTT (120Qs) (FL-HD flies). Using various aggregate-specific antibodies (Ko et al., 2018, Chongtham et al., 2021), we did not detect any HTT aggregates in HD flies treated with or without *E. coli* (Supplementary fig. 3). However, we cannot rule out the formation of unstable oligomers or novel conformations, which may not bind to antibodies used in these studies.

**Figure 1.**
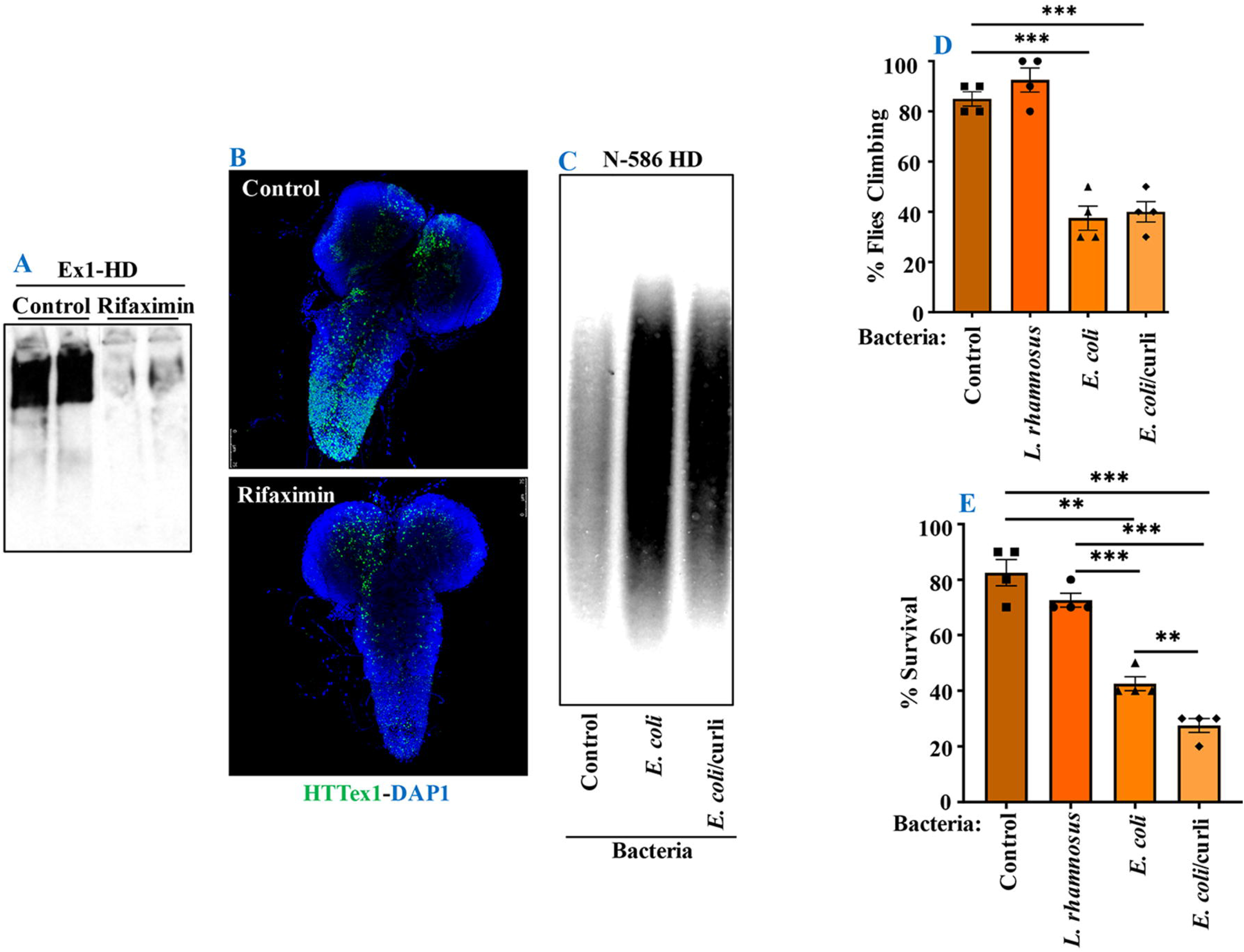
Microbiota modulate HTT aggregation, locomotor behavior and lifespan of *Drosophila* HD models. (A) Western blot detection of mutant HTTex1 aggregates in the lysates of untreated or rifaximin-treated Ex1-HD larvae, using anti-HTT (PHP1) antibody. Each lane represents 10 pooled larvae (N=10) isolated from a different fly vial. (B) Representative confocal images of brains (N=3) from the Ex1-HD larvae treated with rifaximin and immunolabelled with anti-HTT (PHP1, green). DAPI (blue) was used to stain the nuclei. (C) SDD-AGE and WB analysis of lysates (50 μg each) derived from the N-586 HD flies untreated (control), or colonized with *E. coli or E. coli* expressing curli. PHP1 antibody was used to detect HTT aggregates. For each condition 10 larvae were pooled together for analysis (N=10). (D) N-586 HD flies were fed curli-producing/deficient *E. coli or* L*. rhamnosus.* A climbing assay was performed at day 20 after eclosion as described in M&M. Data are reported as mean ± SEM and were analyzed by one-way ANOVA with Tukey’s post hoc test. ***p<0.001, n = 6 groups of 10 flies. (E) The percentage of flies surviving at day 20 was calculated and plotted for each experimental condition. Data are represented as mean ± SEM and were analyzed by one-way ANOVA with Tukey’s post hoc test. ***p<0.001; **p<0.01, n = 4 groups of 10 flies.

### 3.2. Gut Bacteria contribute to immobility and death of HD flies

Abnormal motor behavior is a cardinal symptom of HD. Gut bacteria have been implicated in the locomotion of *Drosophila* (Shretter et al., 2018). Given that *E. coli* promotes HTT aggregation, we asked whether it may alter the mobility of HD flies. The N-586 HD flies do not develop mutant HTT-mediated motor defects, however, colonization with *E. coli* significantly impairs their climbing ability. Colonization with *Lactobacillus rhamnosus* (JB-1 strain) *(L. rhamnosus),* a gram-positive human probiotic has no negative effects (Fig. 1D). Moreover, *E. coli* does not alter the mobility of non-transgenic flies or those expressing WT HTT (Supplementary fig. 4A). These findings suggest that *E. coli* may promote the toxicity of N-586 HTT fragment, which manifests as motor defects in the transgenic flies. *E. coli* also shortens the lifespan of the N-586 HD flies; by day 20 ∼ 50% of the flies die and the rest remain immobile and expire within few days (Fig. 1E). Curli-producing *E. coli* appears more toxic than the isogenic mutant strain indicating that the bacterial amyloids may affect the mortality of N586-HD flies independent of HTT aggregation and locomotive defects (Fig. 1E). Seeding of mutant HTT species isolated from the brains of HD *Drosophila* models has been linked to disease progression and toxicity (Ast et al., 2018). We recently reported that seeding-competent HTTex1 species produce neurotoxic assemblies in human neurons and in neuronal lysates and seeding is further induced by proteinase-K treatment of seeds (Chongtham et al., 2021). To determine whether *E. coli* colonization enhances the seeding competency of mutant HTT in the *E. coli*-treated N-586 HD flies, we performed seeding assays with or without PK treatment. Using equivalent amount of brains lysates as seeds, we find that the seeding activity of mutant HTT treated with PK is enhanced in flies colonized with *E. coli +/-* curli (Supplementary fig. 5). The elevated seeding activity is consistent with immobility defects and enhanced mortality of *E. coli-*treated N586-HD flies and suggest that gut bacteria may influence the production and/or the stability of seeding-competent and potentially neurotoxic HTT species.

Flies expressing full-length human mutant HTT (FL-HD) develop progressive motor defects when compared to those expressing non-pathogenic HTT (25Qs) (Fig. 2A). To examine the role of gut bacteria on the mobility of FL-HD, we treated ∼3 days old adult flies with rifaximin or penicillin-streptomycin and evaluated their climbing ability at different intervals. Antibiotics (ABX) treatment ameliorates the motor defects of FL-HD flies (Figs. 2A, 4A). Conversely, colonization of FL-HD flies with *E. coli* exacerbates the climbing defects but has no effects on the non-transgenic flies or those expressing full-length WT HTT (Fig. 2B, Supplementary fig. 4). Colonization of FL-HD or control flies with *L. rhamnosus* does not alter their climbing abilities. (Fig. 2B, Supplementary fig. 4). Notably, feeding excess *Acetobacter senegalensis* (*A. senegalensis*), a commensal gram-negative bacterium of our *Drosophila* stock also accelerates the motor defects of FL-HD flies (Fig. 2B). The similar debilitating climbing defects induced by two *E. coli* strains and *A. senegalensis* are consistent with a pathogenic role of gram-negative bacteria in the motor behavior of HD flies. *E. coli* colonization also significantly reduces the lifespan of FL-HD flies and similar to the N-586 HD model, the curli-producing strain is more pathogenic (Fig. 2C). Immunostaining the gut of *E. coli*-fed HD flies (FL-HD or the N-586 HD) shows accumulation of *E. coli* in the midgut (Fig. 2D, Supplementary fig. 6), which contains enterocytes and enteric neurons regulating host-microorganism interactions (Dutta et al., 2015, Capo et al., 2019). The localized accumulation of *E. coli* in the midgut is noteworthy as it confirms spatial colonization and a niche, which may participate in gut-brain communications in HD flies.

**Figure 2.**
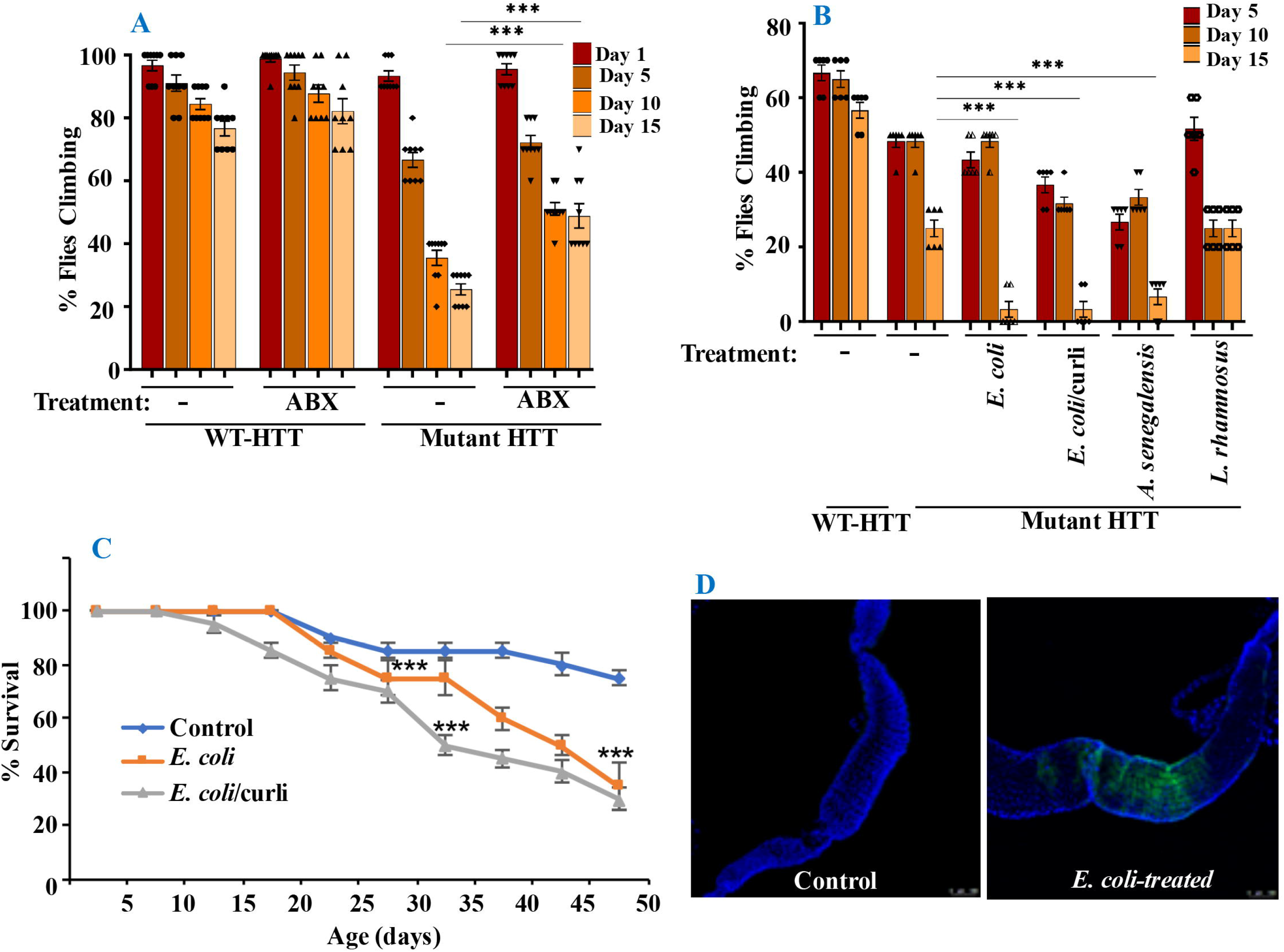
Gut bacteria regulate the motor function and lifespan of FL-HD *Drosophila*. (A) Newly eclosed FL-HD flies or controls expressing full length HTT with 25Qs (WT-HTT) were treated with penicillin-streptomycin (ABX) for 15 days to eliminate gut bacteria. A climbing assay was performed at days 1, 5, 10 and 15 post-treatment. The data are graphed as mean ± SEM, two-way ANOVA with Tukey’s multiple-comparisons test. ***p<0.001, n = 6 groups of 10 flies. (B) Bacteria-deficient adult FL-HD flies were fed different strains of *E. coli*, *A. senegalensis, L. rhamnosus* for 15 days. Locomotive behavior for each condition was quantified as in part A. The data are reported as mean ± SEM, two-way ANOVA with Tukey’s multiple-comparisons test. ***p<0.001; **p<0.01, n = 6 groups of 10 flies. Part C shows the survival curve for FL-HD flies treated with two *E. coli* strains. The percent of flies that survived over time was calculated at different time points. The data are represented as mean ± SEM, two-way ANOVA with Tukey’s multiple-comparisons test. ***p<0.001, n = 6 groups of 10 flies. (D). Immunostaining of the GI tract of control or *E. coli* (curli) treated FL-HD flies demonstrating bacterial colonization of fly gut. A monoclonal antibody generated to *E. coli* was used for detection. DAPI was used to stain the nuclei.

### 3.3. Crocin ameliorates HD phenotypes in Drosophila models

Gut-based regulation of mutant HTT neurotoxicity is a potential therapeutic target. Given that rifaximin treatment of Ex1-HD flies reduces the buildup of neurotoxic HTTex1 assemblies in neurons (Fig. 1), we examined whether gut-based natural compounds might have a similar effect. We focused on anti-inflammatory compounds since inflammation occurs early in pre-manifest HD patients and is recognized as a modifier of HD pathogenesis (Politis et al., 2015). Moreover, inflammation is a prominent outcome of dysbiosis linked to neurodegeneration (Sampson et al., 2016, Goyal et al., 2021). Therefore, we generated Ex1-HD larvae on foods containing luteolin, a flavone which we have found to inhibit the IKKβ-dependent aggregation of HTTex1 in human neurons (Khoshnan et al., 2017), or crocin, a carotenoid abundant in the medicinal plant *Crocus sativus* (saffron), also known for its anti-inflammatory properties and inhibiting the aggregation of amyloidogenic proteins like α-synuclein (Inoue et al., 2018, Shafiee et al., 2018). We find that similar to antibiotics (rifaximin or penicillin-streptomycin) treatment with luteolin or crocin reduces the aggregation of HTTex1 in the nervous system of Ex1-HD larvae (Figs. 1A and B, 3A and B, Supplementary fig. 7). Crocin or antibiotics also enhance the eclosion efficiency of Ex1-HD flies (Fig. 3C) further reinforcing gut-mediated regulation of HTTex1 aggregation, neurotoxicity and survival.

**Figure 3.**
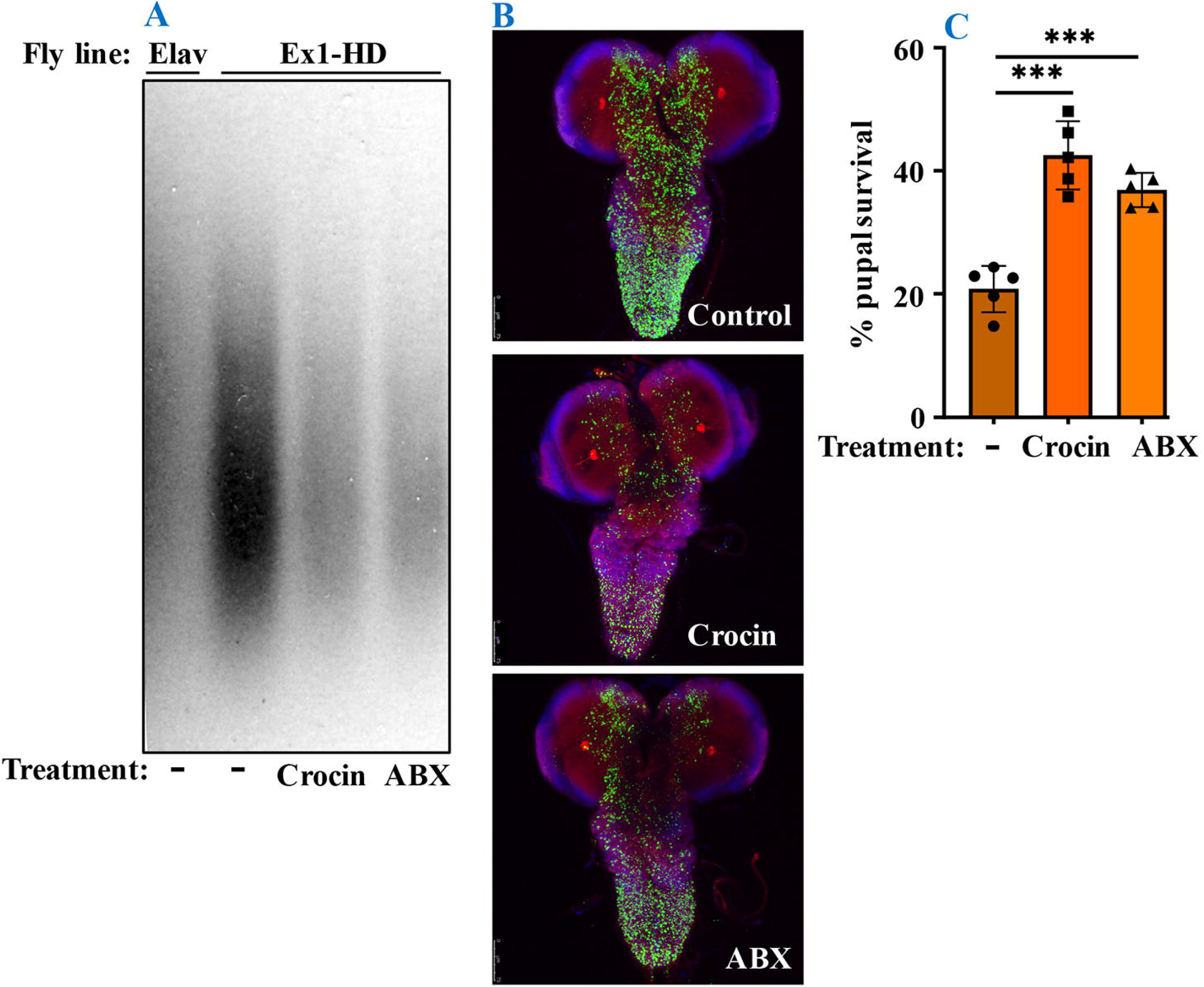
Crocin and antibiotics reduce HTTex1 aggregation and mortality. (A) SDD-AGE and WB analysis of lysates generated from the Ex1-HD larvae treated with crocin or penicillin/streptomycin (ABX). PHP1 antibody was used to detect aggregates. Elav-Gal4 larvae were used as a negative control. For each condition, ten larvae (n=10) were pooled and analyzed. (B) Representative confocal images of brains from Ex1-HD larvae untreated (Control) and treated with crocin or ABX, and immunolabelled with PHP1 (green), anti-Elav antibody (red) and DAPI (blue). Part C shows the percent of pupal survival of Ex1-HD larvae treated with crocin or ABX. The data are represented as mean ± SEM, one-way ANOVA with Tukey’s post hoc test. n = 5 independent crosses for each condition, ***p<0.001.

Notably, addition of crocin, rifaximin or luteolin to fly food also ameliorates the motor defects of FL-HD flies with crocin being the most potent compound (Fig. 4A). Crocin also improves the severe motor defects of FL-HD flies colonized with *E. coli* (Fig. 4B). In survival assays, crocin extends the lifespan of FL-HD flies and those treated with *E. coli +/-* curli (Fig. 4C and D, respectively). Mechanistically, we find that crocin treatment blocks the colonization and accumulation *E. coli* in the flies’ gut (Fig. 4E, Supplementary fig. 8). Crocin does not inhibit the growth of E*. coli* in culture thus, its inhibitory effects on gut colonization may be due to the induction of flies’ anti-microbial defense pathways and/or altering the abundance and/or physiology of gut microbiota.

**Figure 4.**
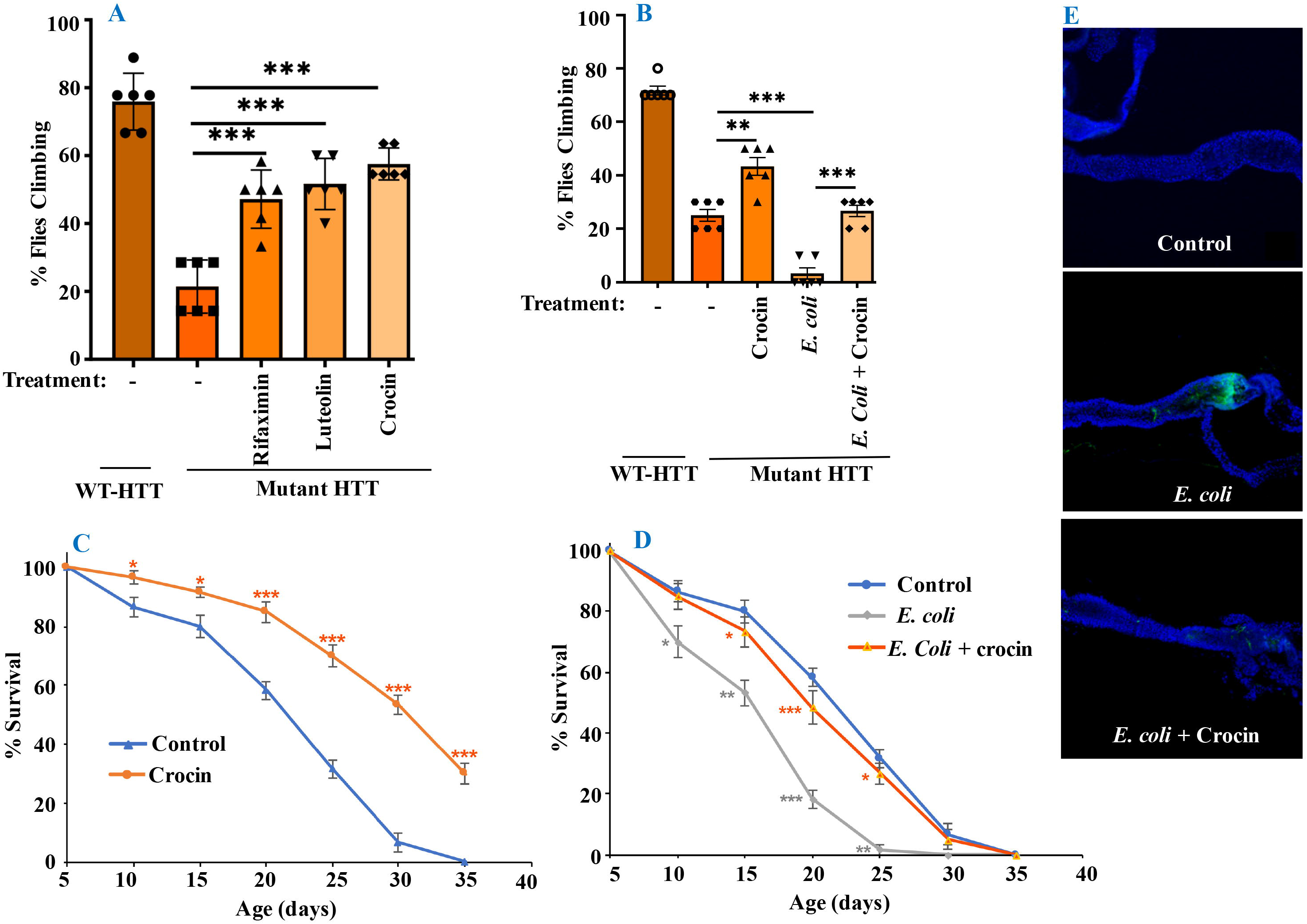
Crocin ameliorates *E. coli*-induced motor defects and mortality in FL-HD flies. (A) Freshly eclosed FL-HD flies were treated with rifaximin, luteolin and crocin for 15 days and their climbing ability was evaluated. Data are reported as mean ± SEM and were analyzed by one-way ANOVA with Tukey’s post hoc test. **p<0.01; *p<0.05, n = 6 groups of 10 flies. (B) Climbing assay was performed to monitor the motor function of flies colonized with crocin, *E. coli* or *E. coli* plus crocin. Untreated FL-HD flies were used as control. Data are represented as mean ± SEM and were analyzed by one-way ANOVA with Tukey’s post hoc test. ***p<0.01; **p<0.01, n = 6 groups of 10 flies. Part C and D show the percentage of FL-HD flies, which survived over time (days) under different treatments. Temperature was elevated to 25 ℃ to accommodate *E. coli* growth. The data are represented as mean ± SEM, two-way ANOVA with Tukey’s multiple-comparisons test. ***p<0.01; **p<0.01; *p<0.05, n = 6 groups of 10 flies. (E) Representative confocal images of the GI tract of untreated (control) FL-HD flies or those treated with *E. coli* (curli) or *E. coli* plus crocin for 15 days showing bacterial colonization and suppression by crocin treatment. Immunostaining was performed using a monoclonal antibody reactive to E*. coli.* DAPI was used to stain the nuclei.

### 3.4. Crocin modulates the composition of gut bacteria in flies

We performed culture assays to examine any potential dysbiosis in the gut of FL-HD flies. Notably, compared to flies expressing WT HTT, FL-HD flies begin to show dysbiosis by day 5 post eclosion exemplified by lower colony-forming units (CFUs) of *Lactobacilli* but elevated CFUs of *Acetobacter* (Supplementary Fig. 9). These findings support an expanded polyQ-mediated disruption in the composition of gut bacteria in FL-HD flies. To gain better insights and examine the impact of crocin on gut bacteria, we performed 16S rRNA sequencing. Our *Drosophila* stocks including those with human HTT transgenes predominantly harbor several *Lactobacilli* strains and *Acetobacter senegalensis (A. senegalensis)* species. Notably, *Lactobacilli* and *Acetobacter* show temporal fluctuation in abundance independent of any transgenes, which may be related to flies’ physiology and nutritional demand (Schretter et al., 2018, Storelli et al., 2018, Henriques et al., 2020, Yamauchi et al., 2020, Ankrah et al., 2021) (Fig. 5, Supplementary fig. 10A). Flies expressing HTT transgenes (WT or mutant) differ with respect to the relative abundance of *Lactobacilli* and *A. senegalensis* when compared to non-transgenic flies (Da-Gal4) (Fig. 5 A-C, Supplementary fig. 10A). Consistent with culture findings, FL-HD flies harbor elevated levels of *Acetobacter* and changes in the abundance of various *Lactobacilli* species when compared to normal flies or those expressing WT HTT (Fig. 5B and C, Supplementary fig. 10A). Interestingly, elevation of *Acetobacter* has been linked to induced mortality in *Drosophila* (Obata et al., 2018). Indeed, FL-HD flies fed excess *A. senegalensis* develop severe motor defects, which is similar in magnitude to those induced by *E. coli* (Fig. 2B). These data are consistent with the induction of dysbiosis by mutant HTT and a potential link to disease progression. Crocin treatment alters the abundance of *Acetotbacter* and *Lactobacillus* in all tested fly lines independent of HTT transgene expression (Fig. 5D-F, Supplementary fig. 10B). Crocin reduces the elevated levels of *Acetobacter* but increases the abundance of *Lactobacillus-NA* (not annotated in data bank) in 5-day old FL-HD flies, however, the effects do not persist (Fig. 5F, Supplementary fig. 10B). We predict that crocin may alter the signaling pathways, which regulate the abundance and/or the physiology of *Lactobacilli* and *Acetobacter* species based on the flies’ nutrients demand (Storelli et al., 2018, Yamauchi et al., 2020). Such systemic and complex metabolic changes may translate into protection observed in the crocin-treated HD flies and remains to be investigated at a molecular level.

**Fig. 5.**
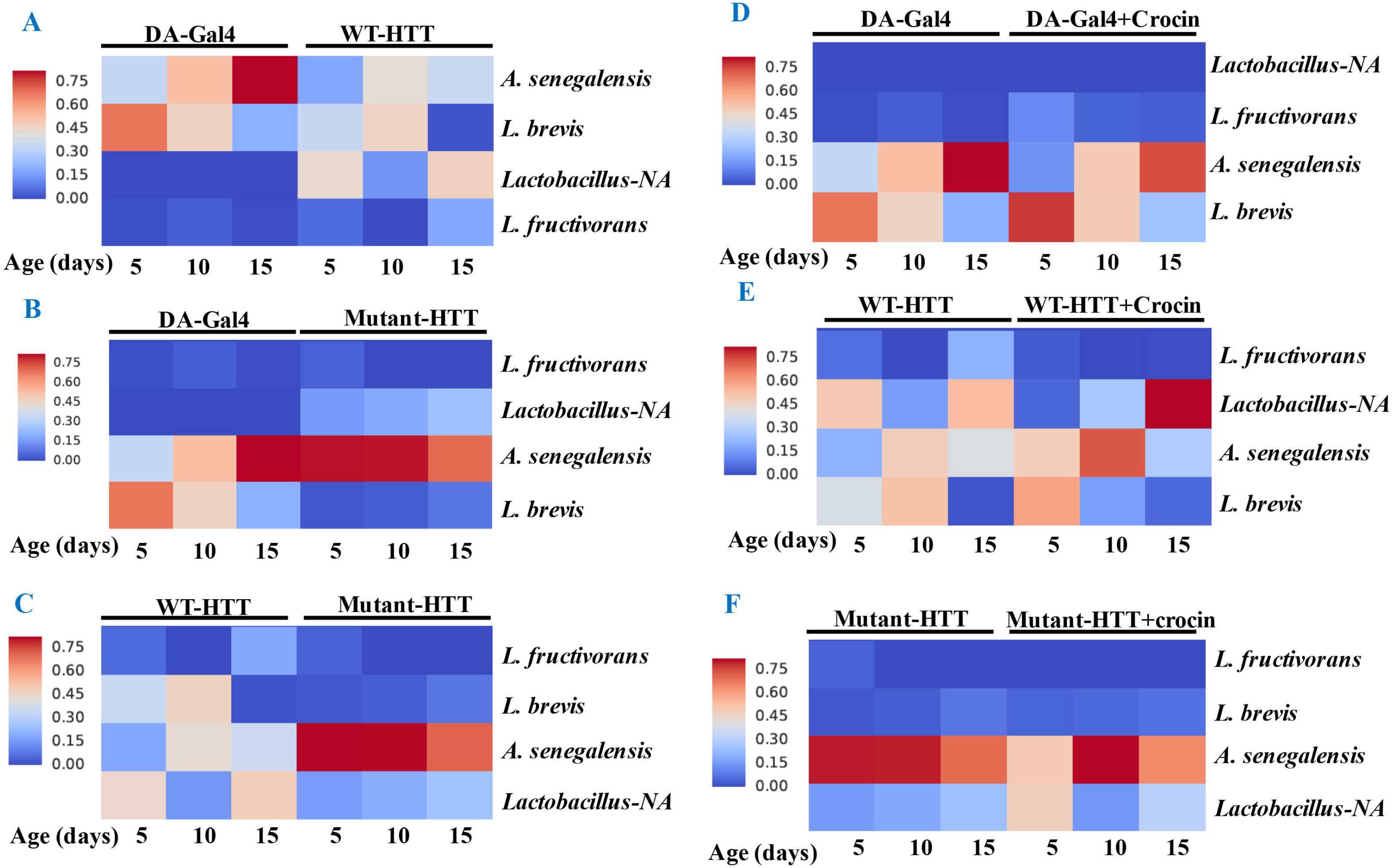
Relative abundance of *Lactobacilli* strains and *A. senegalensis* in the gut of HD flies by 16S rRNA gene sequence analysis. (A) Comparative analysis of DA-Gal4 flies with those expressing WT-HTT, (B) DA-Gal4 with FL-HD flies and (C) WT-HTT with FL-HD. Parts D-F show the effects of crocin on the abundance of bacteria in DA-Gal4, WT-HTT and FL-HD flies, respectively. All analyses were performed as described in the materials and methods by Zymogen’s computational biologists. *Lactobacillus-NA* strain was not detected in the data base.

## 4. Discussion

Recent studies in HD patients and mouse models support a potential role for gut microbiota in disease progression (Kong et al., 2020, Wasser et al., 2020, Du et al., 2021, Kong et al., 2021). However, how a single commensal or opportunistic gut bacterium may influence the pathogenesis of HD remains challenging considering the complexity of gut microbiota and redundancies in functions attributed by various resident microorganisms in mammals. We took advantage of the simplicity of microbiota in *Drosophila* to uncover how bacteria may affect the progression of HD. We provide evidence that gut bacteria promote mutant HTT aggregation, impair locomotion and reduce the lifespan of transgenic HD flies. More specifically, we have identified gram-negative *Acetobacter,* a commensal fly bacterium, and *E. coli* a human pathobiont, as modifiers of HD pathogenesis. As a proof of concept, we have further demonstrated that editing the gut environment of HD flies by a gut-specific antibiotic rifaximin, which is approved to treat dysbiosis in Hepatic Encephalopathy patients (Bureau et al., 20201), or small nutraceuticals such as luteolin or crocin delays HD progression. These studies are prelude to future mechanistic investigations of microbiota-brain interactions in HD and identification of pathways, which may serve as gut-based therapeutic targets.

A hallmark of HD is the accumulation of huntingtin protein aggregates, some of which may be toxic. Our results indicate that elimination of gut bacteria by antibiotics in the Ex1-HD flies significantly reduces the aggregation of amyloidogenic N-terminal HTTex1. The findings are further reinforced by the induction of mutant HTT aggregation by two strains of *E. coli* in the N-586 HD flies. Interestingly, *E. coli* colonization elevated the seeding activity of mutant HTT, which is linked to severity of symptoms in HD animal models and neurotoxicity in human neurons (Ast et al., 2018., Chongtham et al., 2021). The notion that gut bacteria may alter the structure and function of mutant HTT assemblies is a novel finding, which requires biophysical investigations to identify any potential neurotoxic conformations (Ko et al., 2018, Chongtham et al., 2021). Recent studies indicate that gram-negative bacteria promote the aggregation of polyQ peptides and impair the motility of *Caenorhabditis elegans* by disrupting proteostasis in neurons, muscle and intestinal cells (Walker et al., 2021). Our studies are consistent with these findings and underscore the regulatory role of gut bacteria in promoting the misfolding and aggregation of amyloidogenic HTT fragments and the production of neurotoxic assemblies. Surprisingly, several bacterial species have been detected in the brain tissues of HD patients. Among them are members of *Enterobacteriaceae* family, which include *E. coli* and other gram-negative bacteria (Alonso et al., 2019). How bacteria gain access to the brains of HD patients and whether they affect the proteostasis of HTT remain unknown. Our results however, indicate that bacteria such as *E. coli* or potentially *Acetobacter* may signal the aggregation of HTT from the gut. Notably, antibiotic treatments of Ex1-HD flies reduced the aggregation of HTTex1 selectively expressed in neurons (Figs. 1A and B, 3A and B) thus, supporting an indirect role for the gut microbiota in the misfolding and aggregation of amyloidogenic HTT fragments in the nervous system. *E. coli* expressing the functional amyloids curli in the gut has been implicated in α-synuclein aggregation and accelerated PD pathology in the brain of rodents (Chen et al., 2016, Sampson et al., 2020). Our data did not show a selective impact of curli-producing *E. coli* on HTT aggregation and impaired locomotion of HD flies but curli had a negative effect on lifespan and promoted HTTex1 aggregation in mammalian tissue culture (Figs. 1 and 2, 4, Supplementary fig. 2). These findings suggest that curli amyloids may need close contact with HTT or require other factors to promote its aggregation and is potentially unable to enter *Drosophila* cells. Moreover, the negative impact of curli on the survival of HD flies may be independent of HTT aggregation. Notably, *E. coli* and elevated *Acetobacter* (a fly commensal) produced similar debilitating effects on locomotion in HD flies, which is consistent with gram-negative bacteria being modifiers of HD symptoms. A recent study demonstrated a correlation between the abundance of the gram-negative *Bilophila* species and elevated inflammatory cytokine response in HD patients (Du et al., 2021). Our findings along with studies on the microbiota of HD patients and mammalian models support the potential involvement of specific clade of gut bacteria regulating HD progression and severity (Kong et al., 2020, Stan et al., 2020, Wasser et al., 2020, Kong et al., 2021, Du et al., 2021). Given the abundance of inflammogens such as LPS or flagellin in the outer membranes of gram-negative bacteria, and the hypersensitivity of immune cells of HD patients to LPS (Trager et al., 2014), it will be interesting to investigate whether disruptions of gut-brain networks by molecules in the outer membranes of gram-negative bacteria contribute to the pathogenesis of HD.

The ability to modify HD pathology from the gut is therapeutically useful and attractive. Inclusion of crocin in the diet suppressed HTT aggregation, ameliorated locomotive defects and increased the survival of HD flies including those colonized with *E. coli.* The effects of crocin on the diversity of gut bacteria in *Drosophila* were difficult to dissect due to few commensal species. However, crocin treatment altered the abundance of *Lactobacillus* and *Acetobacter* in the young FL-HD flies (Fig. 5F), and prevented *E. coli* colonization in the gut (Fig. 4E, Supplementary fig. 8). In rodents, crocin modifies gut microbiota and prevents neurodegeneration in cerebral ischemia models (Zhang et al., 2019), and mitigates glucocorticoids-induced dysbiosis in mice by increasing the alpha diversity of gut bacteria and subsequent normalization of aberrant lipid metabolism (Xie et al., 2019). Crocin and its major byproduct crocetin do not penetrate the gut epithelium in mammals suggesting that its protective effects are potentially induced by modification of gut physiology and/or its microbiota (Zhang et al., 2019, Xie et al., 2019). Given that alpha diversity of gut bacteria is decreased in HD patients and in HD mice (Kong et al., 2020, Stan et al., 2020, Wasser et al., 2020, Du et al., 2021), the application of crocin as a promoter of bacterial diversity in HD models with complex microbiota may provide useful knowledge on specific species, which regulate HD progression. In addition to identifying gut biomarkers, such studies may also produce novel therapeutic targets for HD.

In summary, our data support a link between gut bacteria and pathogenesis in transgenic *Drosophila* models of HD and highlight the involvement of gram-negative bacteria in regulating the proteostasis and toxicity of mutant HTT. The neurotoxic phenotypes such as protein aggregation, motor defects and mortality induced by *E. coli* are useful biomarkers to investigate the gut-brain circuits at a cellular and molecular level. The *Drosophila* GI tract shares anatomical and functional similarities to mammalian equivalent (Apidianakis and Rahme, 2011). The midgut where *E. coli* colonizes (Figs. 2 and 4) is enriched in numerous cell types including the enteroendocrine (EE) cells, which perform similar tasks as in human GI cells and are nodes for gut-brain networks implicated in neurodegeneration (Chandra et al., 2017). Future knowledge on the transcriptome and proteomics of EE cells in HD *Drosophila* models may identify some of the earliest events, which may trigger the neurotoxicity of mutant HTT in the nervous system. The inclusion of crocin as an inhibitory agent of *E. coli* toxicity may also identify overlapping pathways, potentially functioning in the opposite directions, or compensatory circuits beneficial to HD. Crocin has a long history of medicinal use and has produced favorable outcomes in clinical and preclinical trials in CNS disorders including AD, PD, ischemia, mood disorders and LPS-induced cognitive decline (Rajaei et al., 2016, Zhang et al., 2018, Haeri et al., 2019, Zhang et al., 2019, Azmand and Rajaei 2021). The therapeutic benefits of crocin in subjects with depression have been attributed to the induction of neurortophins expression including brain-derived neurotropic factor (BDNF) (Ghasemi et al., 2015, Moghadam et al., 2021). This property of crocin may prove useful for HD considering the role of HTT in BDNF expression and transport, biological activities that are diminished in neurons expressing mutant HTT (Zuccato and Cattaneo, 2014). The anti-oxidative stress properties of crocin are also noteworthy and may contribute to its therapeutic effects observed in our studies (Cerda-Bernad et al., 2020). Oxidative stress disrupts numerous signaling pathways in HD, which may include inflammatory circuits and genes recently identified in transgenic *Drosophila* models of HD (Kumar and Ratan, 2016, Al-Ramahi et al., 2018, Onur et al., 2021). These features and the gut-modifying properties of crocin make it or its derivatives attractive therapeutic candidates for HD.

## Funding

Funding was provided by awards to AK by Huntington’s Disease Society of America (HDSA) and Hereditary Disease Foundation (HDF).

## Authors’ contributions

AK conceived the idea wrote the manuscript and acquired the funding. AC, JHY, TC, NDA, and AH performed the experiments. AK, AC, and JHY analyzed the data. All authors have read and approved the paper.

## Declaration of Competing Interest

Authors declare no competing interest.

## Acknowledgements

We are grateful to Dr. Segelski at Stanford for providing the *E. coli* strains used in these studies, Dr. J. Lawrence Marsh for providing the HD fly lines and Yelim Lee for technical assistance.

## Notes

### Competing Interest Statement

The authors have declared no competing interest.

### Summary of Updates

The text and figures have been edited and more data included in the supplementary section.

## References

Ast, A., et al., 2018. mHTT Seeding Activity: A Marker of Disease Progression and Neurotoxicity in Models of Huntington’s Disease. Mol. Cell. 71, 675–688.

Abdel-Haq R, Schlachetzki JCM, Glass CK, Mazmanian SK., 2019. Microbiome-microglia connections via the gut-brain axis. J Exp Med 216, 41–59.

Alonso R, Pisa D, Carrasco L., 2019. Brain Microbiota in Huntington’s Disease Patients. Front Microbiol 10, 2622.

Al-Ramahi I, Lu B, Di Paola S, Pang K, de Haro M, Peluso I, Gallego-Flores T, Malik NT, et al., 2018. High-Throughput Functional Analysis Distinguishes Pathogenic, Nonpathogenic, and Compensatory Transcriptional Changes in Neurodegeneration. Cell Syst 7, 28–40 e24.

Ankrah NYD, Barker BE, Song J, Wu C, McMullen JG, 2nd, Douglas AE., 2021. Predicted Metabolic Function of the Gut Microbiota of Drosophila melanogaster. mSystems 6, e01369-20.

Apidianakis Y, Rahme LG., 2011. Drosophila melanogaster as a model for human intestinal infection and pathology. Dis Model Mech 4, 21–30.

Azmand MJ, Rajaei Z., 2021. Effects of crocin on spatial or aversive learning and memory impairments induced by lipopolysaccharide in rats. Avicenna J Phytomed 11, 79–90.

Barbaro BA, Lukacsovich T, Agrawal N, Burke J, Bornemann DJ, Purcell JM, Worthge SA, Caricasole A, et al., 2015. Comparative study of naturally occurring huntingtin fragments in Drosophila points to exon 1 as the most pathogenic species in Huntington’s disease. Hum Mol Genet 24, 913–925.

Bates GP, Dorsey R, Gusella JF, Hayden MR, Kay C, Leavitt BR, Nance M, Ross CA, et al., 2015. Huntington disease. Nat Rev Dis Primers 1:15005.

Björkqvist M, Wild EJ, Thiele J, Silvestroni A, Andre R, Lahiri N, Raibon E, Lee RV, et al., 2008. A novel pathogenic pathway of immune activation detectable before clinical onset in Huntington’s disease. J Exp Med 205, 1869–1877.

Bureau C, Thabut D, Jezequel C, Archambeaud I, D’Alteroche L, Dharancy S, Borentain P, Oberti F, et al., 2021. The Use of Rifaximin in the Prevention of Overt Hepatic Encephalopathy After Transjugular Intrahepatic Portosystemic Shunt : A Randomized Controlled Trial. Ann Intern Med 174, 633–640.

Capo F, Wilson A, Di Cara F., 2019. The Intestine of Drosophila melanogaster: An Emerging Versatile Model System to Study Intestinal Epithelial Homeostasis and Host-Microbial Interactions in Humans. Microorganisms 7.

Caporaso, J., Kuczynski, J., Stombaugh, J. et al., 2010. QIIME allows analysis of high-throughput community sequencing data. Nat Methods 7, 335–336.

Cerda-Bernad D, Valero-Cases E, Pastor JJ, Frutos MJ., 2020. Saffron bioactives crocin, crocetin and safranal: effect on oxidative stress and mechanisms of action. Crit Rev Food Sci Nutr:1–18.

Chandra R, Hiniker A, Kuo YM, Nussbaum RL, Liddle RA., 2017. alpha-Synuclein in gut endocrine cells and its implications for Parkinson’s disease. JCI Insight 2.

Chen SG, Stribinskis V, Rane MJ, Demuth DR, Gozal E, Roberts AM, Jagadapillai R, Liu R, et al., 2016. Exposure to the Functional Bacterial Amyloid Protein Curli Enhances Alpha-Synuclein Aggregation in Aged Fischer 344 Rats and Caenorhabditis elegans. Sci Rep 6, 34477.

Chongtham A, Bornemann DJ, Barbaro BA, Lukacsovich T, Agrawal N, Syed A, Worthge S, Purcell J, et al.,2020. Effects of flanking sequences and cellular context on subcellular behavior and pathology of mutant HTT. Hum Mol Genet 29, 674–688.

Chongtham A., et al., 2021. Amplification of neurotoxic HTTex1 assemblies in human neurons. Neurobiol Dis. 159, 105517.

DiFiglia M, Sapp E, Chase KO, Davies SW, Bates GP, Vonsattel JP, Aronin N., 1997. Aggregation of huntingtin in neuronal intranuclear inclusions and dystrophic neurites in brain. Science 277,1990–1993.

Douglas AE, 2018. The Drosophila model for microbiome research. Lab Anim (NY) 47,157–164.

Du G, Dong W, Yang Q, Yu X, Ma J, Gu W, Huang Y., 2021. Altered Gut Microbiota Related to Inflammatory Responses in Patients With Huntington’s Disease. Front Immunol 11,603594.

Dutta D, Dobson AJ, Houtz PL, Glasser C, Revah J, Korzelius J, Patel PH, Edgar BA, et al., 2015. Regional Cell-Specific Transcriptome Mapping Reveals Regulatory Complexity in the Adult Drosophila Midgut. Cell Rep 12, 346–358.

Ghasemi T, Abnous K, Vahdati F, Mehri S, Razavi BM, Hosseinzadeh H., 2015.Antidepressant Effect of Crocus sativus Aqueous Extract and its Effect on CREB, BDNF, and VGF Transcript and Protein Levels in Rat Hippocampus. Drug Res (Stuttg) 65, 337–343.

Ghosh R, Tabrizi SJ., 2018. Clinical Features of Huntington’s Disease. Adv Exp Med Biol 1049, 1–28.

Goyal D, Ali SA, Singh RK., 2021. Emerging role of gut microbiota in modulation of neuroinflammation and neurodegeneration with emphasis on Alzheimer’s disease. Prog Neuropsychopharmacol Biol Psychiatry 106, 110112.

Haeri P, Mohammadipour A, Heidari Z, Ebrahimzadeh-Bideskan A., 2019. Neuroprotective effect of crocin on substantia nigra in MPTP-induced Parkinson’s disease model of mice. Anat Sci Int 94, 119–127.

Halfmann R, Lindquist S., 2008. Screening for amyloid aggregation by Semi-Denaturing Detergent-Agarose Gel Electrophoresis. J Vis Exp. 17, 838.

Hanson MA, Lemaitre B., 2020 New insights on Drosophila antimicrobial peptide function in host defense and beyond. Curr Opin Immunol. 62, 22–30.

Hatfield I, Harvey I, Yates ER, Redd JR, Reiter LT, Bridges D., 2015. The role of TORC1 in muscle development in Drosophila. Sci Rep 5, 9676.

Henriques SF, Dhakan DB, Serra L, Francisco AP, Carvalho-Santos Z, Baltazar C, Elias AP, Anjos M, et al., 2020. Metabolic cross-feeding in imbalanced diets allows gut microbes to improve reproduction and alter host behaviour. Nat Commun 11, 4236.

Hsiao EY, McBride SW, Hsien S, Sharon G, Hyde ER, McCue T, Codelli JA, Chow J, et al., 2013. Microbiota modulate behavioral and physiological abnormalities associated with neurodevelopmental disorders. Cell 155, 1451–1463.

Inoue E, Shimizu Y, Masui R, Hayakawa T, Tsubonoya T, Hori S, Sudoh K., 2018. Effects of saffron and its constituents, crocin-1, crocin-2, and crocetin on alpha-synuclein fibrils. J Nat Med 72, 274–279.

Keshavarzian A, Green SJ, Engen PA, Voigt RM, Naqib A, Forsyth CB, Mutlu E, Shannon KM., 2015. Colonic bacterial composition in Parkinson’s disease. Mov Disord 30, 1351–1360.

Khoshnan A, Sabbaugh A, Calamini B, Marinero SA, Dunn DE, Yoo JH, Ko J, Lo DC, et al., 2017. IKKbeta and mutant huntingtin interactions regulate the expression of IL-34: implications for microglial-mediated neurodegeneration in HD. Hum Mol Genet 26, 4267–4277.

Khoshnan A, Ko J, Watkin EE, Paige LA, Reinhart PH, Patterson PH., 2004.Activation of the IkappaB kinase complex and nuclear factor-kappaB contributes to mutant huntingtin neurotoxicity. J Neurosci 24, 7999–8008.

Ko, J et al., 2018. Identification of distinct conformations associated with monomers and fibril assemblies of mutant huntingtin. Hum. Mol. Genet. 27, 2330–2343.

Kong G, Cao KL, Judd LM, Li S, Renoir T, Hannan AJ., 2020. Microbiome profiling reveals gut dysbiosis in a transgenic mouse model of Huntington’s disease. Neurobiol Dis 135, 104268.

Kong G, Ellul S, Narayana VK, Kanojia K, Ha HTT, Li S, Renoir T, Cao KL, et al., 2021. An integrated metagenomics and metabolomics approach implicates the microbiota-gut-brain axis in the pathogenesis of Huntington’s disease. Neurobiol Dis 148,105199.

Kumar A, Ratan RR., 2016. Oxidative Stress and Huntington’s Disease: The Good, The Bad, and The Ugly. J Huntingtons Dis 5,217–237.

Li W, Wu X, Hu X, Wang T, Liang S, Duan Y, Jin F, Qin B., 2017. Structural changes of gut microbiota in Parkinson’s disease and its correlation with clinical features. Sci China Life Sci 60, 1223–1233.

Liu Z, Wang X, Yu Y, Li X, Wang T, Jiang H, Ren Q, Jiao Y, et al., 2008. A Drosophila model for LRRK2-linked parkinsonism. Proc Natl Acad Sci U S A 105, 2693–2698.

Marizzoni M, Cattaneo A, Mirabelli P, Festari C, Lopizzo N, Nicolosi V, Mombelli E, Mazzelli M, et al., 2020. Short-Chain Fatty Acids and Lipopolysaccharide as Mediators Between Gut Dysbiosis and Amyloid Pathology in Alzheimer’s Disease. J Alzheimers Dis 78:683–697.

Moghadam BH, Bagheri R, Roozbeh B, Ashtary-Larky D, Gaeini AA, Dutheil F, Wong A., 2021. Impact of saffron (Crocus Sativus Linn) supplementation and resistance training on markers implicated in depression and happiness levels in untrained young males. Physiol Behav 233:113352.

Morais LH, Schreiber HLt, Mazmanian SK., 2021.The gut microbiota-brain axis in behaviour and brain disorders. Nat Rev Microbiol 19, 241–255.

Obata F, Fons CO, Gould AP., 2018. Early-life exposure to low-dose oxidants can increase longevity via microbiome remodelling in Drosophila. Nat Commun 9, 975.

Onur TS, Laitman A, Zhao H, Keyho R, Kim H, Wang J, Mair M, Wang H, Li L, Perez A, de Haro M, Wan YW, Allen G, Lu B, Al-Ramahi I, Liu Z, Botas J., 2021. Downregulation of glial genes involved in synaptic function mitigates Huntington’s disease pathogenesis. Elife. 10, e64564.

Perez-Pardo P, Dodiya HB, Engen PA, Forsyth CB, Huschens AM, Shaikh M, Voigt RM, Naqib A, et al., 2019. Role of TLR4 in the gut-brain axis in Parkinson’s disease: a translational study from men to mice. Gut 68, 829–843.

Politis M, Lahiri N, Niccolini F, Su P, Wu K, Giannetti P, Scahill RI, Turkheimer FE, et al., 2015. Increased central microglial activation associated with peripheral cytokine levels in premanifest Huntington’s disease gene carriers. Neurobiol Dis 83, 115–121.

Rajaei Z, Hosseini M, Alaei H., 2016. Effects of crocin on brain oxidative damage and aversive memory in a 6-OHDA model of Parkinson’s disease. Arq Neuropsiquiatr. 74, 723–729.

Reichhardt C, Jacobson AN, Maher MC, Uang J, McCrate OA, Eckart M, Cegelski L., 2015. Congo Red Interactions with Curli-Producing E. coli and Native Curli Amyloid Fibers. PLoS One 10, e0140388.

Sampson TR, Debelius JW, Thron T, Janssen S, Shastri GG, Ilhan ZE, Challis C, Schretter CE, et al., 2016.Gut Microbiota Regulate Motor Deficits and Neuroinflammation in a Model of Parkinson’s Disease. Cell. 167, 1469–1480.e12.

Sampson TR, Challis C, Jain N, Moiseyenko A, Ladinsky MS, Shastri GG, Thron T, Needham BD, et al., 2020. A gut bacterial amyloid promotes alpha-synuclein aggregation and motor impairment in mice. Elife 9.

Shafiee M, Arekhi S, Omranzadeh A, Sahebkar A., 2018. Saffron in the treatment of depression, anxiety and other mental disorders: Current evidence and potential mechanisms of action. J Affect Disord 227, 330–337.

Sharon G, Sampson TR, Geschwind DH, Mazmanian SK., 2016. The Central Nervous System and the Gut Microbiome. Cell 167, 915–932.

Scheperjans F, Aho V, Pereira PA, Koskinen K, Paulin L, Pekkonen E, Haapaniemi E, Kaakkola S, et al., 2015. Gut microbiota are related to Parkinson’s disease and clinical phenotype. Mov Disord 30, 350–358.

Schretter CE, Vielmetter J, Bartos I, Marka Z, Marka S, Argade S, Mazmanian SK., 2018. A gut microbial factor modulates locomotor behaviour in Drosophila. Nature 563, 402–406.

Segata N, Izard J, Waldron L, Gevers D, Miropolsky L, Garrett WS, Huttenhower C., 2011. Metagenomic biomarker discovery and explanation. Genome Biol. 2, R60.

Stan TL, Soylu-Kucharz R, Burleigh S, Prykhodko O, Cao L, Franke N, Sjogren M, Haikal C, et al., 2020. Increased intestinal permeability and gut dysbiosis in the R6/2 mouse model of Huntington’s disease. Sci Rep 10, 18270.

Storelli G, Strigini M, Grenier T, Bozonnet L, Schwarzer M, Daniel C, Matos R, Leulier F., 2018. Drosophila Perpetuates Nutritional Mutualism by Promoting the Fitness of Its Intestinal Symbiont Lactobacillus plantarum. Cell Metab 27, 362–377 e368.

Takewaki D, Suda W, Sato W, Takayasu L, Kumar N, Kimura K, Kaga N, Mizuno T, et al., 2020. Alterations of the gut ecological and functional microenvironment in different stages of multiple sclerosis. Proc Natl Acad Sci U S A 117, 22402–22412.

The Huntington’s Disease Collaborative Research Group, 1993. A novel gene containing a trinucleotide repeat that is expanded and unstable on Huntington’s disease chromosomes. Cell. 72, 971–983.

Trager U, Andre R, Lahiri N, Magnusson-Lind A, Weiss A, Grueninger S, McKinnon C, Sirinathsinghji E, et al., 2014. HTT-lowering reverses Huntington’s disease immune dysfunction caused by NFkappaB pathway dysregulation. Brain 137, 819–833.

von Essen MR, Hellem MNN, Vinther-Jensen T, Ammitzboll C, Hansen RH, Hjermind LE, Nielsen TT, Nielsen JE, et al., 2020. Early Intrathecal T Helper 17.1 Cell Activity in Huntington Disease. Ann Neurol 87, 246–255.

Walker AC, Bhargava R, Vaziriyan-Sani AS, Pourciau C, Donahue ET, Dove AS, Gebhardt MJ, Ellward GL, et al., 2021. Colonization of the Caenorhabditis elegans gut with human enteric bacterial pathogens leads to proteostasis disruption that is rescued by butyrate. PLoS Pathog 17, e1009510.

Wasser CI, Mercieca EC, Kong G, Hannan AJ, McKeown SJ, Glikmann-Johnston Y, Stout JC., 2020. Gut dysbiosis in Huntington’s disease: associations among gut microbiota, cognitive performance and clinical outcomes. Brain Commun 2, fcaa110.

Wu SC, Cao ZS, Chang KM, Juang JL., 2017. Intestinal microbial dysbiosis aggravates the progression of Alzheimer’s disease in Drosophila. Nat Commun 8, 24.

Yamauchi T, Oi A, Kosakamoto H, Akuzawa-Tokita Y, Murakami T, Mori H, Miura M, Obata F., 2020. Gut Bacterial Species Distinctively Impact Host Purine Metabolites during Aging in Drosophila. iScience, 101477.

Zeng Q, Shen J, Chen K, Zhou J, Liao Q, Lu K, Yuan J, Bi F., 2020. The alteration of gut microbiome and metabolism in amyotrophic lateral sclerosis patients. Sci Rep 10, 12998.

Zhang L, Previn R, Lu L, Liao RF, Jin Y, Wang RK., 2018. Crocin, a natural product attenuates lipopolysaccharide-induced anxiety and depressive-like behaviors through suppressing NF-kB and NLRP3 signaling pathway. Brain Res Bull 142, 352–359.

Zhang Y, Geng J, Hong Y, Jiao L, Li S, Sun R, Xie Y, Yan C, et al., 2019. Orally Administered Crocin Protects Against Cerebral Ischemia/Reperfusion Injury Through the Metabolic Transformation of Crocetin by Gut Microbiota. Front Pharmacol 10, 440.

Xie X, Xiao Q, Xiong Z, Yu C, Zhou J, Fu Z., 2019. Crocin-I ameliorates the disruption of lipid metabolism and dysbiosis of the gut microbiota induced by chronic corticosterone in mice. Food Funct. 10,6779–6791.

Zuccato C, Cattaneo E., 2014. Huntington’s disease. Handb Exp Pharmacol 220, 357–409.

