## Supplemental methods for "Gut bacteria regulate the pathogenesis of Huntington’s disease in *Drosophila* model"

*PCR amplification bacteria*

Flies were surface sterilized with 70% ethanol for 1 min and washed three times with sterile 1X PBS. Bacterial DNA was extracted from equal number of HTTex1 (25Qs or 103Qs) using stool DNA kit (Omega Bio-tek, Norcross, Georgia) according to the provided procedures. Bacterial DNA was amplified from 5 ng of total DNA from each batch of fly by q-pCR using the universal primer Eub340F: TCCTACGGGAGGCAGCAGT-3' and Eub781R: 5'-GGACTACCAGGGTATCTAATCCTGTT-3') (Nadkarni et al., 2002) in a 7300 real-time PCR system. Data were analyzed comparatively by the formula 2^(-ΔΔCT)^.

*Construction of pLenti6-csgA-6x His Plasmid*

*E. coli* genomic DNA was isolated from ~1 mL of overnight culture using QIAamp DNA Mini Kit (QIAGEN), following the manufacturer’s protocol. C-terminally hexahistidine-tagged CsgA without N-terminal SEC secretion signal sequence (Evans et al. 2015) was amplified via standard PCR method using the following primers: csgA F: 5’-TCAAGGGAATTCACCATGGGTGTTGTTCCTCAGTACGG-3’, csgA R: 5’-TCAACGGGATCCCTAGTGATGATGGTGGTGATGGTACTGATGAGCGGTCGCGTTG-3’. Thermal cycling program used for the PCR is as follows: 94 ºC, 5 min → 33x [94 ºC, 30 s → 58 ºC, 30 s → 72 ºC, 30s] → 72 ºC, 7 min → 4 ºC. The resulting PCR amplicon was gel-purified using QIAquick Gel Extraction kit (QIAGEN) and then assembled into pLenti6/V5-D-TOPO vector (Invitrogen) following the manufacturer’s protocol. The cloned plasmid was subsequently transformed into One Shot TOP10 chemically competent *E. coli* (Invitrogen) and spread on LB +Ampicillin (100 μg/mL) agar plate for selection. Several plasmid clones were isolated using QIAprep Spin Miniprep kit (QIAGEN) and then screened via agarose gel electrophoresis after digesting them with EcoRI and BamHI. The correct clone was sequenced using CMV-F primer (5’-CGCAAATGGGCGGTAGGCGTG-3’) and named pLenti6-V5-D-TOPO::csgA-6x His.

*Co-transfection of HTTEx1 Q103-EGFP and csgA Plasmids in HEK293 Cells*

HEK293 cells were seeded in a 6-well plate with approximately 25 % density and let grow overnight at 37 ºC, 5 % CO_2_. The cells were then transfected using calcium phosphate transfection method. In brief, 0.1 μg FUGW::HTTEx1 Q103-EGFP plasmid and differential amounts of pLenti6-V5-D-TOPO::csgA-6x His plasmid (0, 0.2 μg or 0.3 μg), were added in each snap-cap tube. Betagalactase cDNA cloned in similar vector was used as control. The total amount of DNA in each transfection mixture was normalized using an empty plasmid backbone. 2 M CaCl_2_ was subsequently added (0.12 M final concentration), and the volume of mixture was brought up to 100 μL with ddH_2_O. 100 μL 2x HBS was added next dropwise and part of solution was squirted into the rest by pipetting for aeration. Entire volume of each transfection mixture was added to the cells and incubation was done overnight at 37 ºC, 5 % CO_2_. The transfected cells were subjected to widefield fluorescence microscopy for quantifying HTTEx1 Q103-EGFP aggregates and then harvested for SDS-PAGE and western blotting analyses for HTTEx1 Q103-EGFP (probed with PHP2 antibody, 1:1000 dilution) and CsgA-6x His (probed with mouse anti-His antibody, from Thermo Fisher, 1:5000 dilution).

Evans ML, Chorell E, Taylor JD, Åden J, Götheson A, Li F, Koch M, Sefer L, Matthews SJ, et al., (2015), The bacterial curli system possesses a potent and selective inhibitor of amyloid formation. Mol Cell, 57, 445-55.

Nadkarni, MA, Martin, F.E., Jacques, NA, and Hunter, N (2002). Determination of bacterial load by real-time PCR using a broad-range (universal) probe and primers set. Microbiology 148, 257–266.
