## Supplementary figures and images for "Gut bacteria regulate the pathogenesis of Huntington’s disease in *Drosophila* model"

### Suppl. data 11-23-21.pptx

#### Slide 1
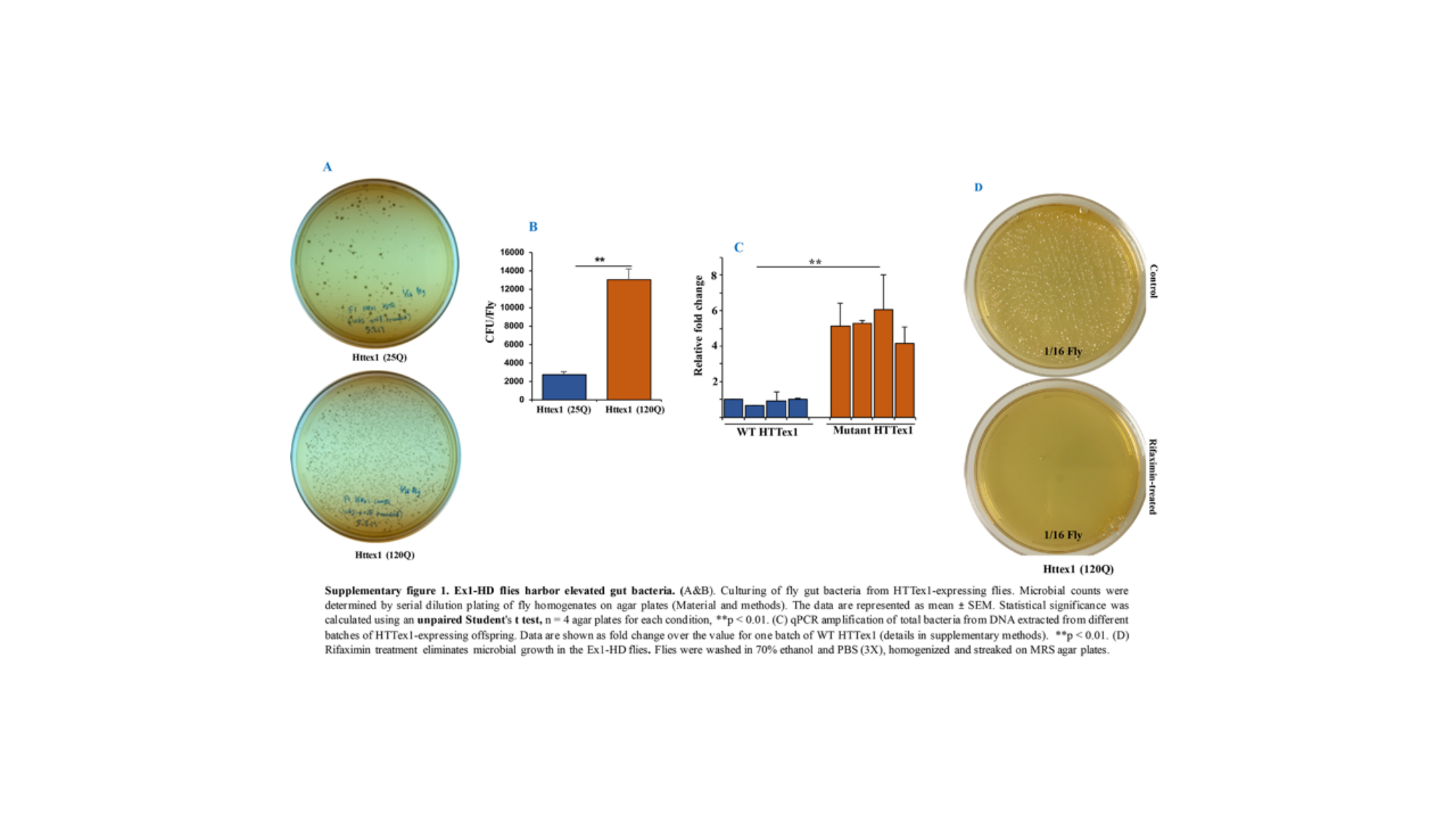

#### Slide 2
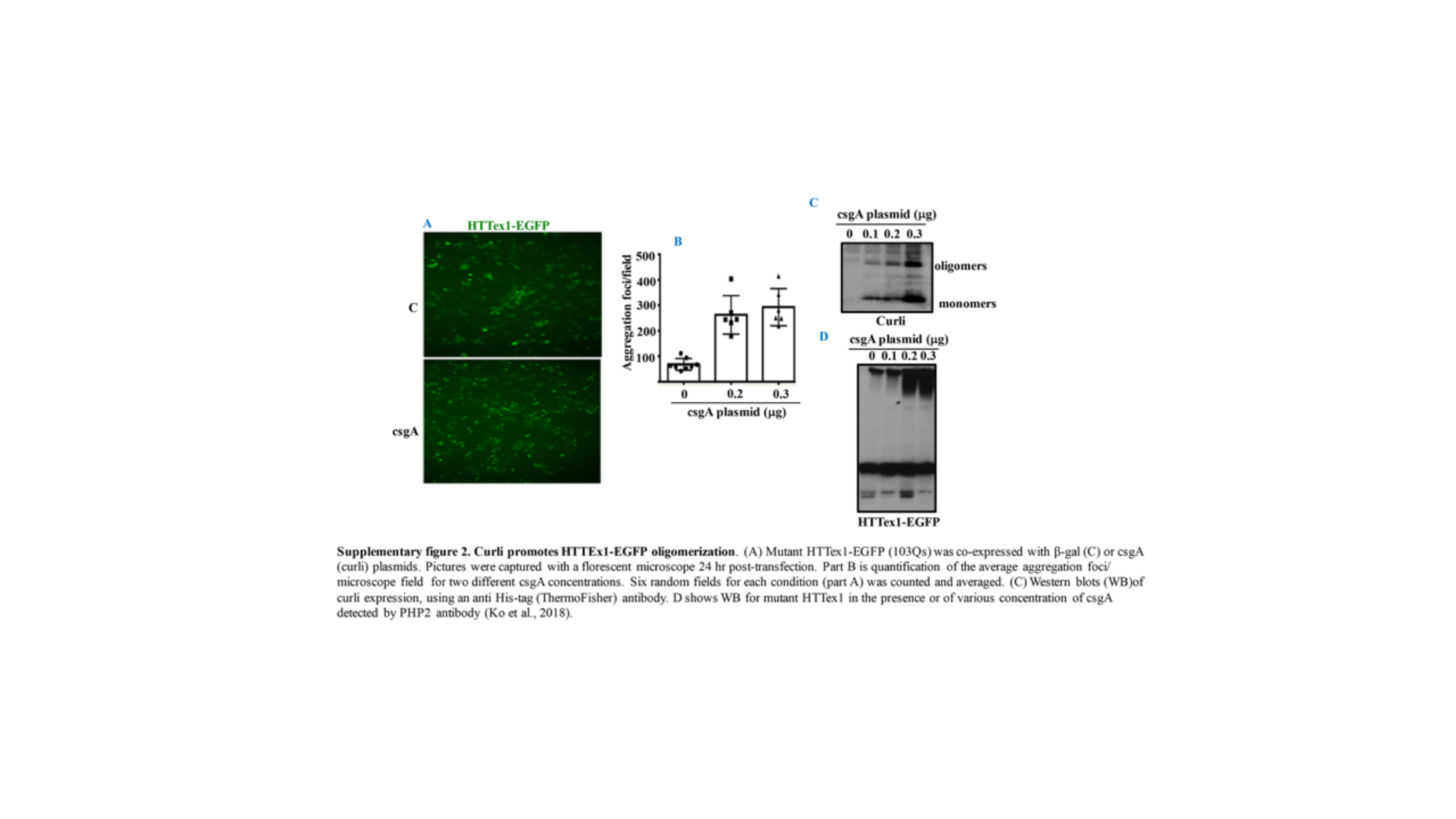

#### Slide 3
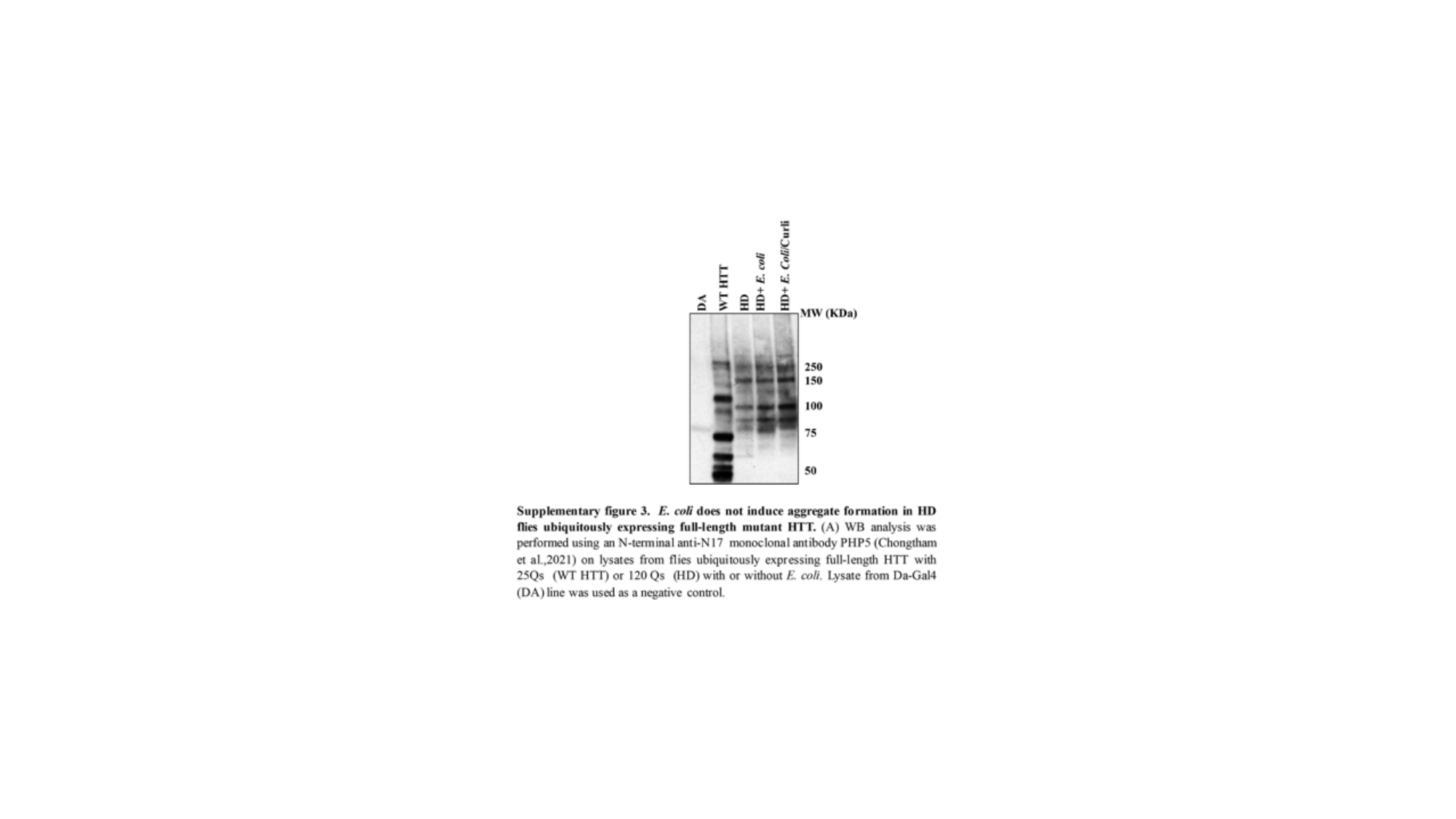

#### Slide 4
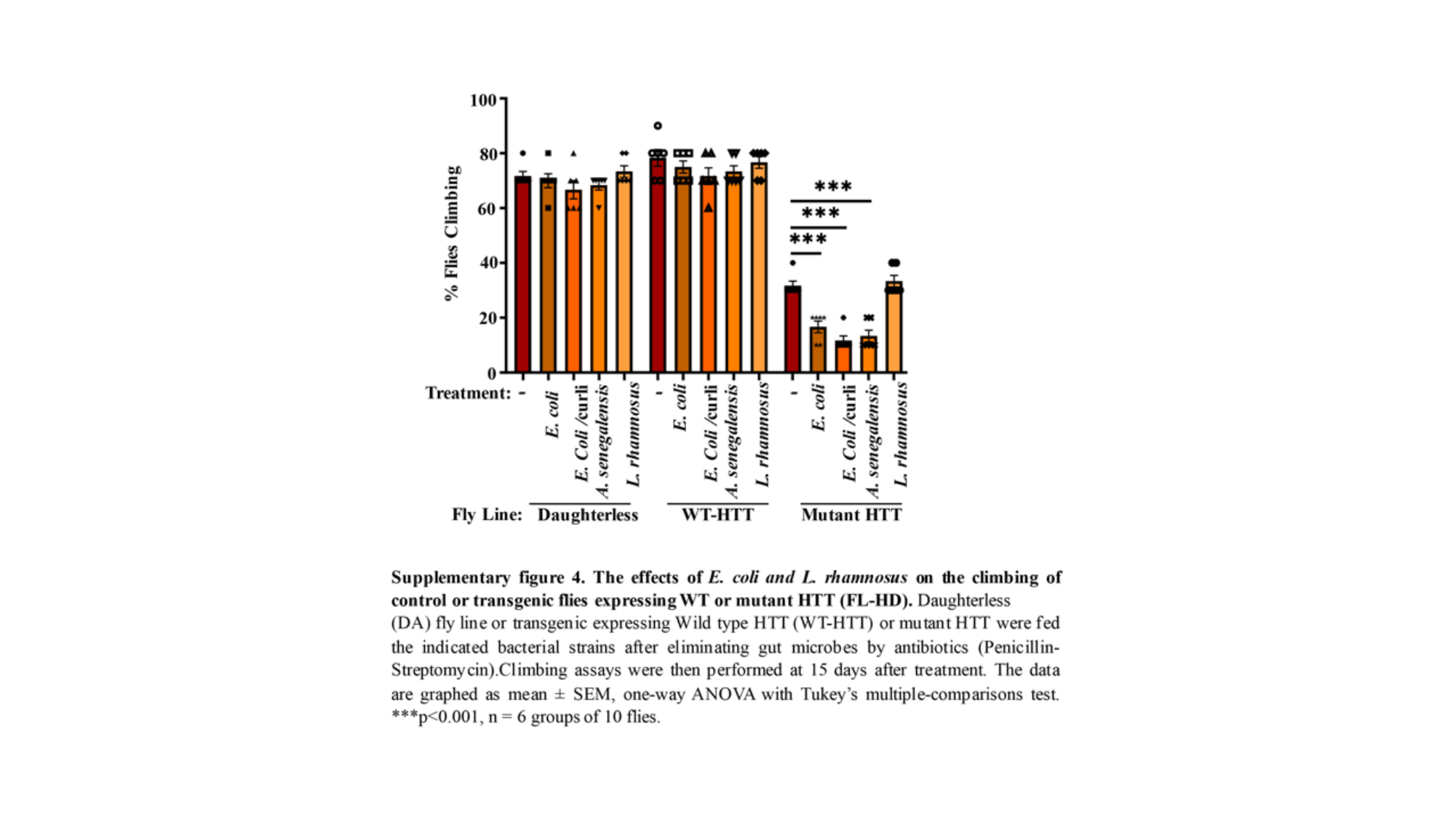

#### Slide 5
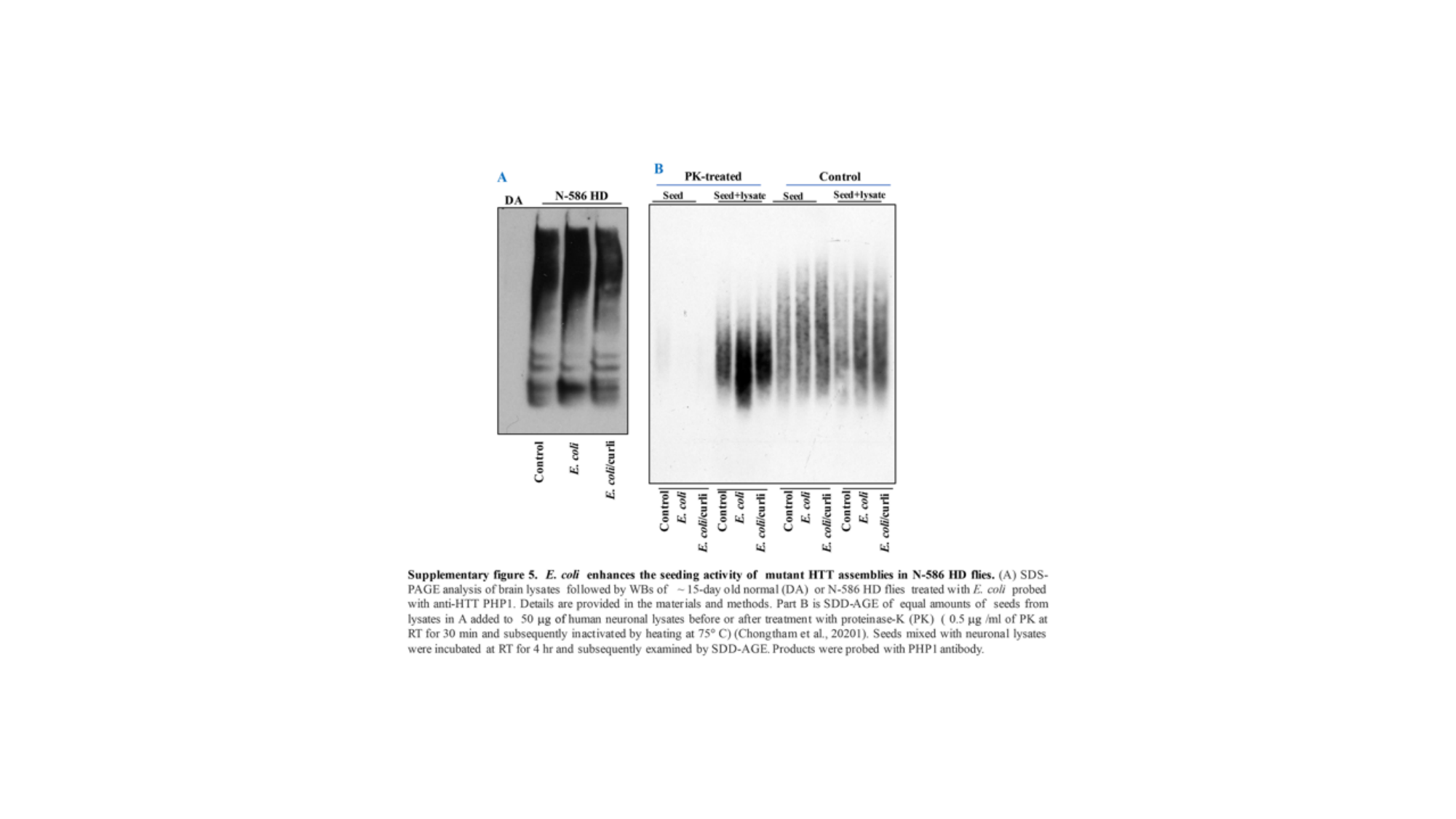

#### Slide 6
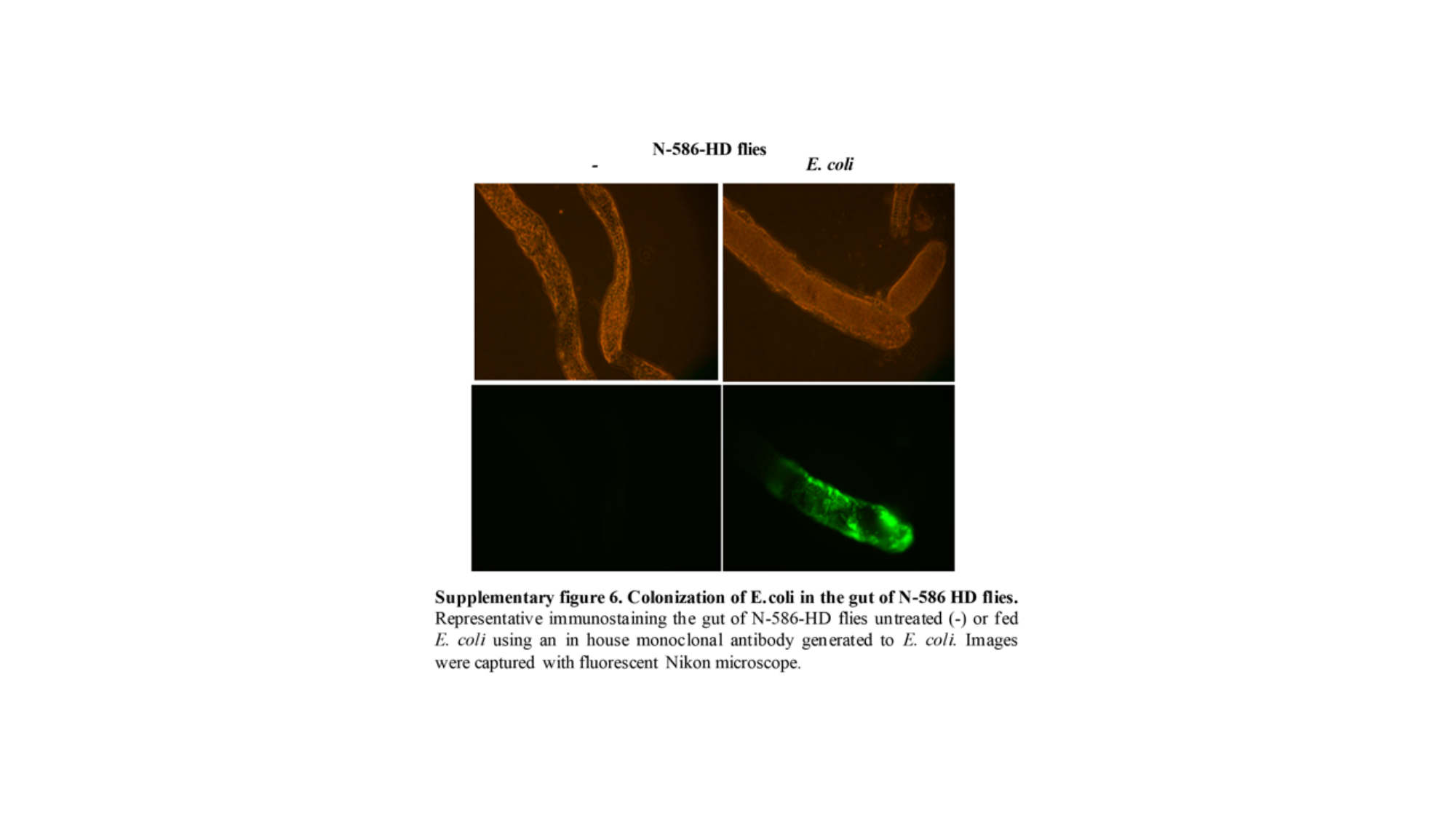

#### Slide 7
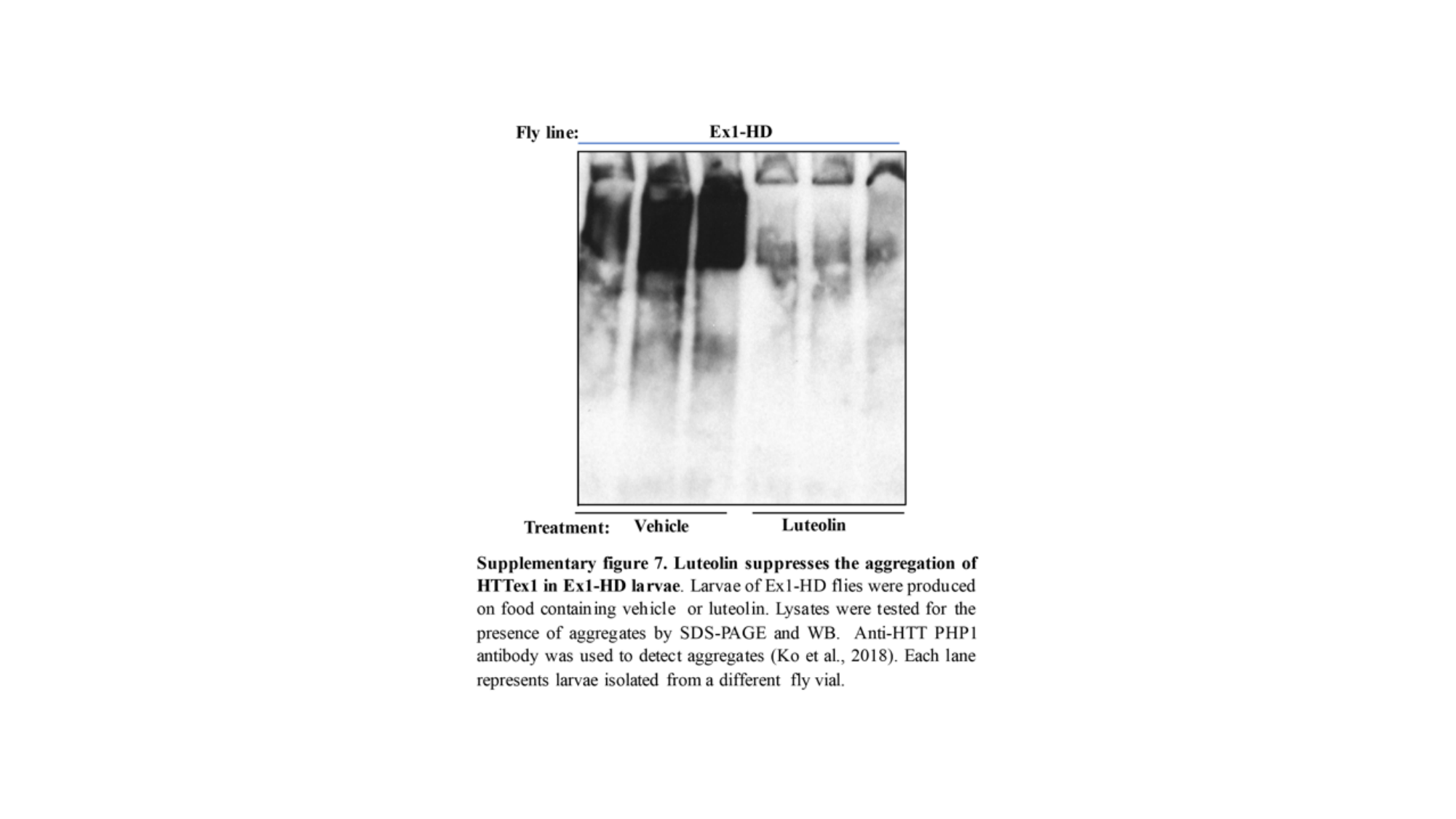

#### Slide 8
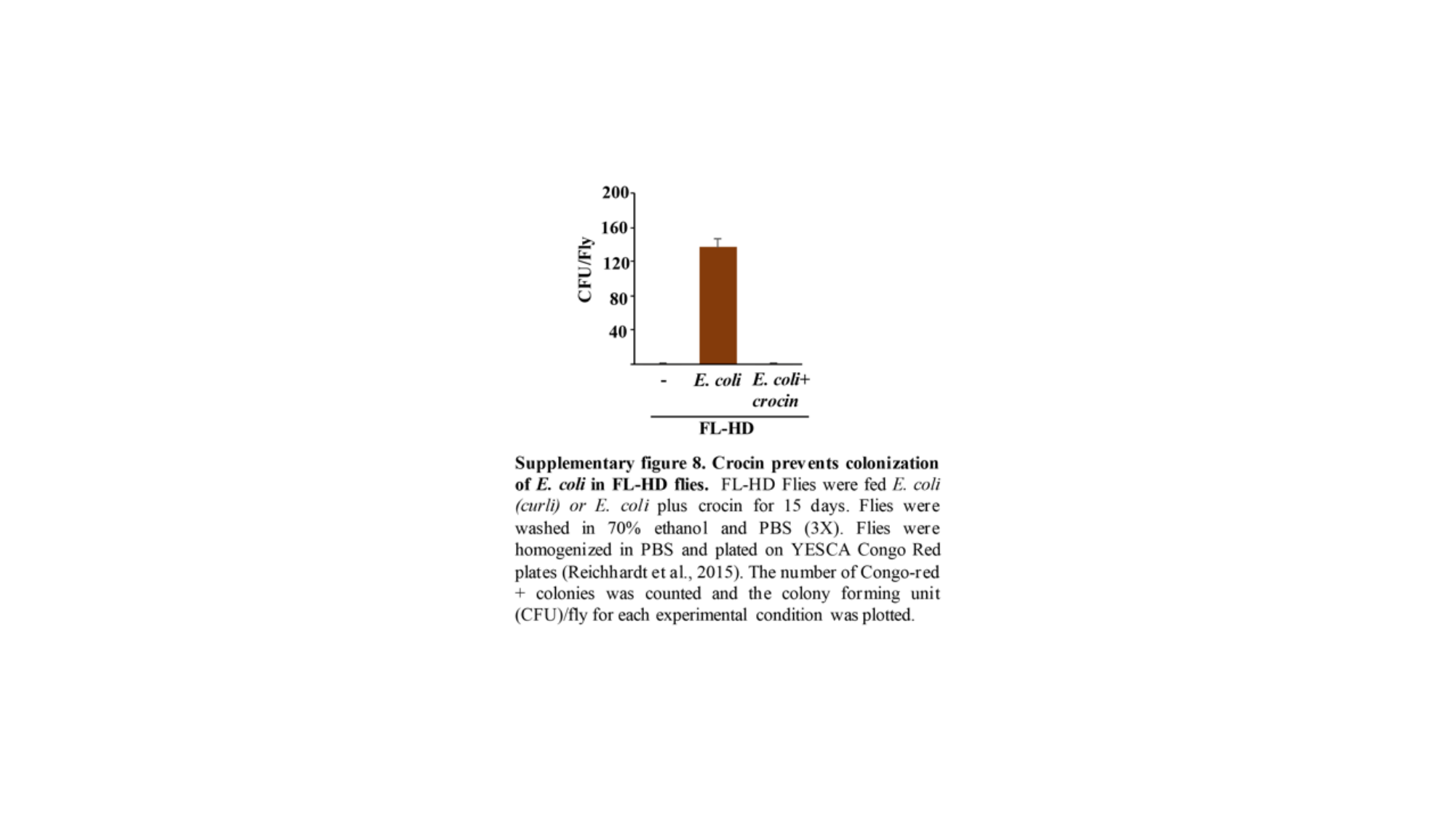

#### Slide 9
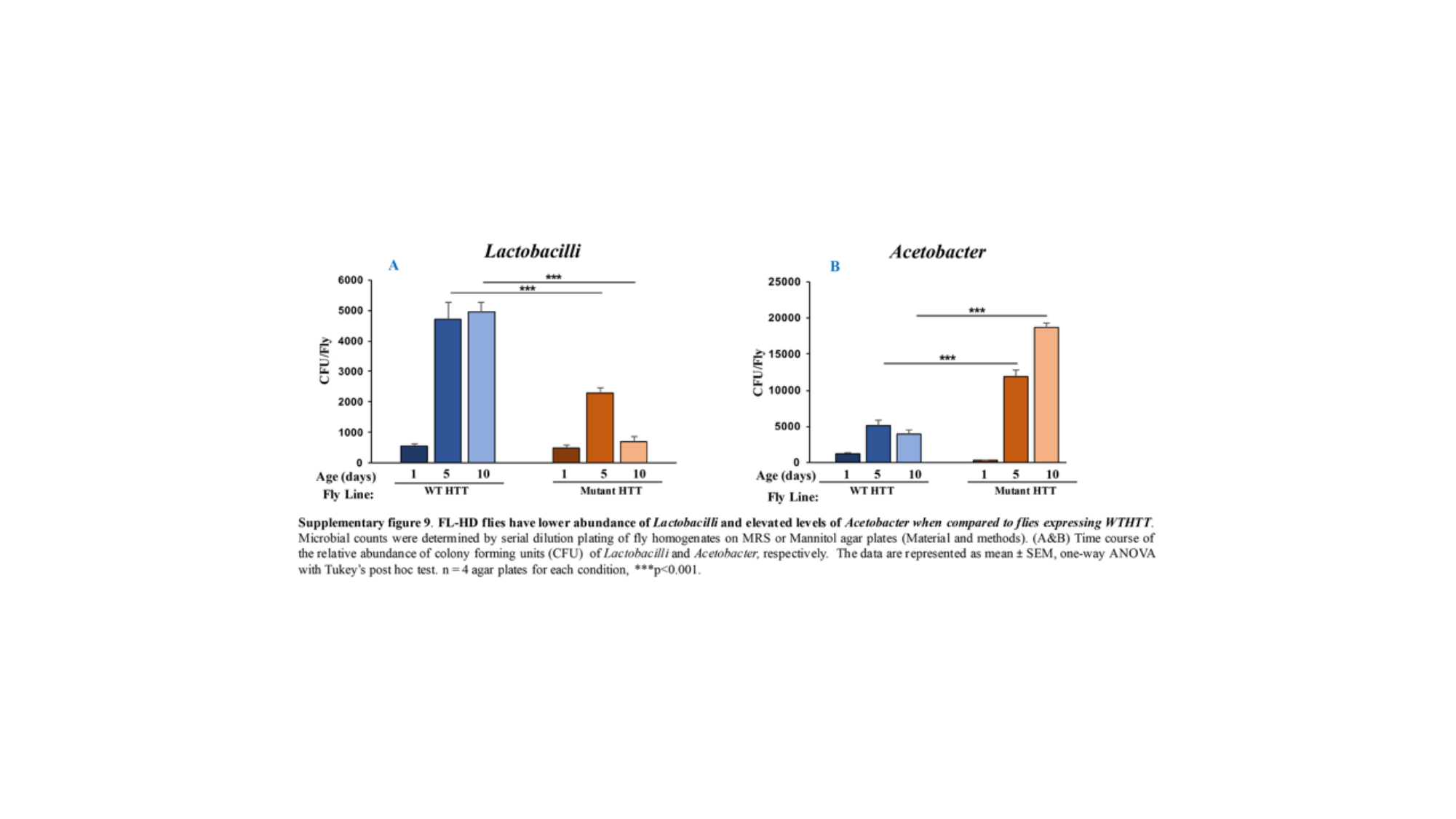

#### Slide 10
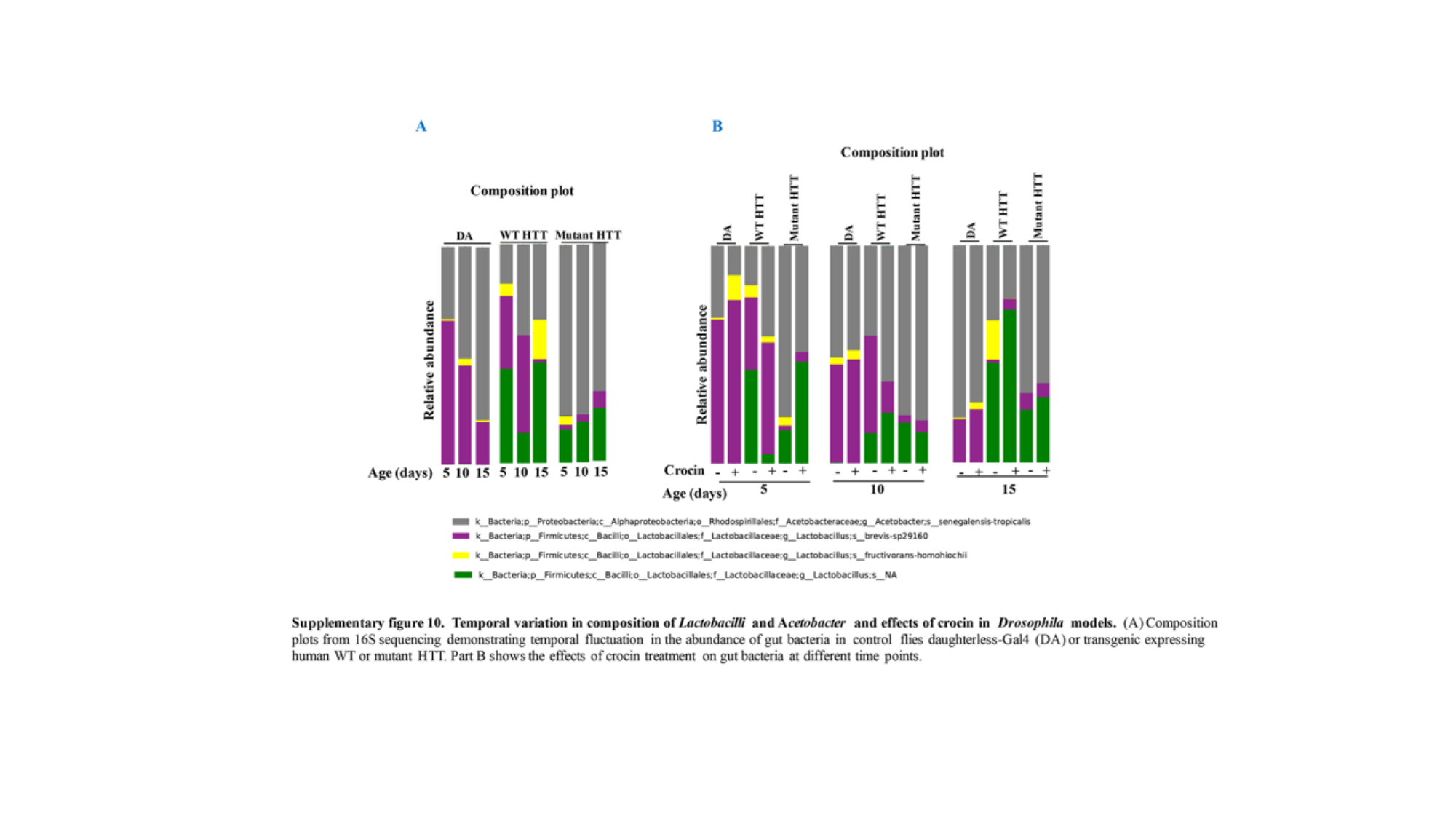
